# Experimental evolution reveals bifunctional genetic solutions to loss of *trpF* in *Salmonella enterica*

**DOI:** 10.64898/2026.01.20.700655

**Authors:** Joakim Näsvall, Hind Abdalaal

**Affiliations:** Dept. of Medical Biochemistry and Microbiology, Uppsala University, Uppsala, Sweden

## Abstract

How new gene functions arise while maintaining ancestral biological roles remains a central question in evolutionary genetics. To investigate genetic solutions to disruption of a biosynthetic pathway without prior genetic bias, we used experimental evolution to study restoration of tryptophan biosynthesis in *Salmonella enterica* strains lacking the *trpF* gene. Populations were founded from bacteria carrying wild-type alleles of all relevant genes in their native genomic and regulatory contexts and evolved under conditions selecting for growth without exogenous tryptophan. Across independent populations, mutations in either *hisA* or *trpA* enabled growth of Δ*trpF* strains in the absence of added tryptophan while retaining sufficient ancestral function to support growth under the same conditions. Whole-genome sequencing and genetic reconstruction showed that these mutant alleles were sufficient to confer the growth-rescue phenotype. Duplication of the target gene was detected in only a single population and showed no evidence of functional divergence. Mutational paths differed between genes: *hisA*-based solutions arose primarily in mutator backgrounds and were associated with stronger trade-offs with ancestral function, whereas *trpA*-based solutions were more frequent and often retained native function. Together, these results demonstrate that bifunctional genetic solutions can arise through point mutations in multiple genes during experimental adaptation, illustrating how gene-level multifunctionality can evolve without gene duplication.

## Introduction

The evolution of new gene functions often involves mutations that alter existing biological roles, creating trade-offs between ancestral and newly acquired activities (Soskine and Tawfik 2010). Such conflicts can be resolved through gene duplication followed by divergence (Ohno 1970; Bergthorsson et al. 2007; Deng et al. 2010; Näsvall et al. 2012). However, although relatively rare in the literature, some genes in nature perform more than one physiologically relevant function, indicating that multifunctionality can also arise and persist without duplication (Mueller et al. 1998; Barona-Gómez and Hodgson 2003). Understanding how genes acquire additional functions while retaining ancestral roles is therefore central to evolutionary genetics, but direct experimental evidence linking specific genetic changes to such outcomes remains limited (Näsvall et al. 2012; Lundin et al. 2020).

One experimental system that has proven particularly informative for studying the evolution of new gene functions involves the histidine and tryptophan biosynthetic pathways in bacteria. In many bacteria, the enzymes HisA and TrpF catalyze chemically related reactions on structurally similar substrates, and both belong to the same family of (αβ)_8_-barrel enzymes, reflecting a shared evolutionary origin (Copley and Bork 2000). Consistent with this relationship, previous work has shown that HisA can acquire TrpF activity through mutation, often accompanied by trade-offs with its native function (Jürgens et al. 2000; Näsvall et al. 2012; Newton et al. 2017; Lundin et al. 2020). In addition, some actinobacterial species lack a dedicated *trpF* gene and instead rely on a single HisA ortholog, PriA, that supports both histidine and tryptophan biosynthesis, demonstrating that physiologically relevant bifunctionality can be maintained in nature (Barona-Gómez and Hodgson 2003; Due et al. 2011; Plach et al. 2016).

Additionally, TrpA, an enzyme that acts later in the tryptophan biosynthetic pathway and is less closely related to TrpF, has also been shown to acquire TrpF activity through directed evolution and rational mutagenesis, indicating that multiple genes can, in principle, provide genetic solutions to loss of *trpF* (Evran et al. 2012). However, most previous experimental studies of TrpF replacement have relied on engineered starting genotypes, altered gene expression, or continuous strong selection, potentially biasing evolutionary outcomes toward particular genes or evolutionary paths (Jürgens et al. 2000; Evran et al. 2012; Näsvall et al. 2012; Lundin et al. 2020). As a result, it remains unclear which genes are most likely to evolve physiologically relevant bifunctionality when selection acts on wild-type genetic backgrounds under minimally manipulated conditions.

To reduce experimental biases that can influence which genetic solutions are observed during adaptation, we designed an experimental evolution framework starting from minimally modified genetic backgrounds. We used *Salmonella enterica* strains lacking trpF but otherwise carrying wild-type alleles of *hisA, trpA*, and all other relevant genes in their native genomic locations and under their native regulation. Because deletion of *trpF* prevents growth in the absence of tryptophan, populations were propagated in medium containing a limiting amount of tryptophan, allowing initial growth of the ancestral genotype while selecting for restoration of tryptophan biosynthesis as the nutrient became depleted. This approach differs from previous studies that relied on engineered starting alleles, altered gene expression, genomic contexts prone to amplification, or targeted mutagenesis, all of which can bias evolutionary trajectories toward particular genes or mechanisms (Jürgens et al. 2000; Evran et al. 2012; Näsvall et al. 2012; Lundin et al. 2020).

Using this framework, we followed the early stages of adaptation across replicate populations and identified the genetic changes that enabled growth of Δ*trpF* strains without added tryptophan. Whole-genome sequencing and reconstruction of evolved alleles were then used to determine which mutations were sufficient to confer the growth-rescue phenotype, allowing us to characterize the genetic basis and functional consequences of the adaptations that emerged.

## Materials and Methods

### Bacterial strains and growth conditions

All bacterial strains used in this study (listed in Table S3) are derivatives of *Salmonella enterica* subsp. *enterica* serovar Typhimurium strain LT2, except for three *Escherichia coli* strains used as PCR templates for amplification of selection cassettes employed in λ Red recombineering. Unless otherwise stated, generalized transduction was performed using phage P22 *HT105*/*1 int*-*201* (Schmieger 1972) or the defective P22-derived artificial gene transfer agent GTA22 described in the Supplementary Methods.

Lysogeny broth (LB; 10 g/L tryptone, 5 g/L yeast extract, 10 g/L NaCl) was used as rich medium, with 15 g/L Bacto agar added for solid medium (LA). Salt-free LB (LB without NaCl) was used for preparing electrocompetent cells. SOC medium (Hanahan 1983) was used for recovery following electroporation. As minimal medium, M9 medium (Miller 1992) was prepared at double strength with respect to M9 salts, glucose, CaCl_2_, and MgCl_2_ (hereafter referred to as 2× M9) to allow growth to higher cell densities. Where indicated, minimal medium was supplemented with tryptophan (0.1 mM or 5 μM), histidine (0.1 mM), and/or guanosine (0.3 mM). Solid M9 plates contained 15 g/L Bacto agar. All cultures were grown at 37 °C except during λ Red recombineering.

Antibiotics were used only for selection of recombinants following genetic manipulations and included trimethoprim (10 mg/L), tetracycline (7.5 mg/L), and chloramphenicol (12.5 mg/L).

### PCR and Sanger sequencing

PCR products for λ Red recombineering and Sanger sequencing were generated using Phusion DNA polymerase (Thermo Fisher). PCR templates consisted of bacterial cell suspensions prepared either from a portion of a single colony or from ∼1 μL of frozen stock culture resuspended in 100 μL ultrapure water. One microliter of suspension was used per 20 μL PCR reaction. Cycling conditions have been described previously (Näsvall et al. 2017; Näsvall 2017; Näsvall 2022).

PCR products were purified and concentrated by polyethylene glycol (PEG) precipitation using a 2× solution containing 24% (w/v) PEG 8000 and 20 mM MgCl_2_. DNA pellets were washed with 70% ethanol, air-dried, and resuspended in ultrapure water. For Sanger sequencing, purified PCR products were mixed with sequencing primers (Table S4) and submitted to Eurofins Genomics (Ebersberg, Germany).

### λ Red recombineering and genetic reconstructions

Genetic markers, deletions, and allelic replacements were constructed using λ Red recombineering as described previously (Yu et al. 2000; Näsvall et al. 2017; Näsvall 2017; Näsvall 2022). Strains carrying the λ Red helper plasmid pSIM5-Tet (Koskiniemi et al. 2011) were grown overnight at 30 °C in salt-free LB supplemented with 0.2% glucose, diluted 1:100, and induced for recombination by incubation at 42 °C for 15 min. Cells were rendered electrocompetent by washing in ice-cold water or 10% glycerol and electroporated with purified PCR products.

Mutations in *hisA* and *trpA* were reconstructed by PCR amplification of the mutated alleles from evolved populations or clones, followed by replacement of a selectable and counter-selectable cassette (*Atox1*; GenBank MN207489) inserted at the target locus. Selection for loss of the cassette was performed on M9 medium supplemented with rhamnose (0.2%), histidine, and tryptophan. In cases where evolved alleles contained multiple mutations, overlapping PCR fragments derived from mutant and wild-type templates were used to reconstruct specific alleles.

Deletions, insertions, and selected point mutations were constructed using DIRex with half-cassettes derived from *Atox1* and *Atox2* (GenBank MN207490), as described previously (Näsvall 2017; Näsvall 2022). All oligonucleotides are listed in Table S4.

### Experimental evolution

Two experimental evolution studies were performed (outlined in Figure 1).

**Figure 1.**
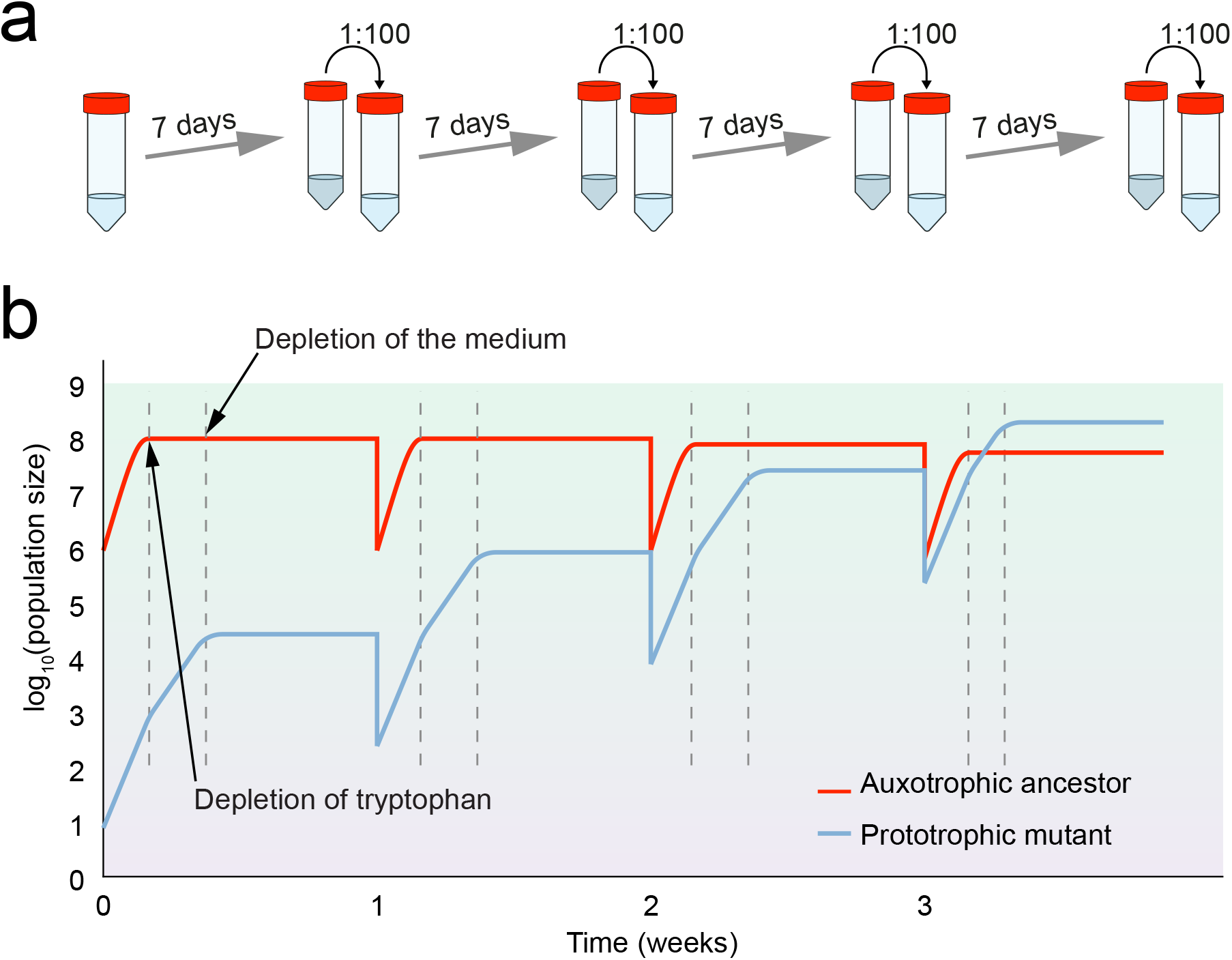
Experimental outline. **(a)** Serial passage experiments with *Salmonella enterica* lacking the *trpF* gene, grown in media with limiting tryptophan. Each 7-day cycle consisted of a short period with unlimited growth (tryptophan present) followed by a longer period without tryptophan (tryptophan-limited selection). Only prototrophic mutants (that had gained TrpF activity) could grow during this phase. **(b)** The expected dynamics during the takeover of a batch culture population by a TrpF+ mutant competing with its TrpF-ancestor in tryptophan-limited medium. Each cycle includes a brief window where the TrpF+ mutant can grow and outcompete the TrpF-ancestor. The TrpF-ancestor is expected to seize growing when tryptophan is depleted while the TrpF+ mutant continues growing until the medium is depleted for some other essential nutrient(s). Initially, the non-growing TrpF-ancestor sets the window by continuing to metabolize and consume resources, thereby limiting the time available for the TrpF+ mutant to grow. However, once the TrpF+ mutant becomes dominant, it will determine the length of the window through its own metabolic activity.

**Experiment 1** was initiated from independent cultures of strain DA52864 (Δ*trpF* Δ*P*_*gal*_ Δ*galE*::*T*_*lux*_). Following overnight growth in medium containing histidine and tryptophan, cultures were diluted 1:100 into 5 mL tryptophan-limited medium (2× M9 + 5 μM tryptophan) and propagated by serial transfer. Cultures were passaged once every seven days for 20 cycles and subsequently twice weekly for an additional 30 cycles, totaling approximately 430 generations. Samples were frozen at −80 °C in 20% DMSO at multiple time points.

**Experiment 2** was initiated from strains DA62207 (Δ*trpF*) and DA64168 (Δ*trpF mutS*), each carrying a defective P22-derived gene transfer agent (see Supplementary Methods: *Construction and use of an artificial Gene Transfer Agent (GTA22)*). Forty-eight populations were passaged weekly by 1:100 dilution in tryptophan-limited medium for 13 cycles. Samples were frozen at −80 °C in 20% glycerol at multiple time points.

### Growth assays and fitness measurements

Growth assays were initiated from overnight cultures grown in 2× M9 supplemented with glucose, histidine, and tryptophan. Cultures were diluted 1:1000 into tryptophan-free or tryptophan-limited medium and transferred to Bioscreen C honeycomb plates. Optical density at 600 nm was recorded every 4 min for up to 7 days at 37 °C with continuous shaking.

Growth rates were estimated by fitting OD data from the exponential phase (OD_600_ ≤ 0.18) to an exponential growth model using GraphPad Prism. Area under the curve (AUC) values were calculated for the first 72 h following blank subtraction to approximate fitness effects. Additional details and considerations including evaporation effects and post-depletion OD fluctuations that can influence AUC estimates are described in the Supplementary Methods.

### Whole-genome re-sequencing

Genomic DNA was extracted from evolving populations using MasterPure DNA purification kits (Epicentre). Libraries for MiSeq sequencing (in-house) were prepared using Nextera XT DNA library preparation and indexing kits (Illumina). Some samples were sequenced by BGI (Beijing, China) using DNBseq technology. Reads were processed and analyzed using CLC Genomics Workbench. Variants were identified by mapping reads to a reference genome containing all known ancestral modifications. Copy number estimates for amplified regions were derived from read depth comparisons as described in the Supplementary Methods.

## Results

### Experimental evolution restores growth of Δ*trpF* strains under intermittent selection

To identify genetic changes that enable growth of *Salmonella enterica* strains lacking *trpF* under minimally biased conditions, we evolved replicate populations under intermittent selection for rescue of tryptophan prototrophy. Because loss of *trpF* prevents growth only after depletion of exogenous tryptophan, cultures were propagated in medium containing a limiting amount of tryptophan. Each growth cycle therefore began with a period during which all cells could grow, followed by a phase in which continued growth depended on genetic changes that restored the ability to synthesize tryptophan (Figure 1).

Eight populations were evolved for approximately 430 generations. By the end of the experiment, three populations exhibited detectable growth after tryptophan depletion (Figure 2b; Table 1). Whole-genome sequencing (WGS) revealed that two of these populations carried multiple amino acid substitutions in *hisA* (Tables 1 and S1). Both populations also harbored mutations previously shown to increase mutation rates (frameshift mutations in *mutH* and *mutS*, respectively; Table S1) and accumulated numerous additional mutations across the genome. The mutator mutations were present at low frequencies after approximately 100 generations (Table S1), preceding the appearance of *hisA* mutations by roughly 30 generations (Table S2).

**Table 1.** Identified mutations conferring TrpF activity.

| Strain # | Population<sup>a</sup> | # Generations<sup>b</sup> | Causative mutation(s)<sup>c</sup> | Identified by<sup>d</sup> |
| --- | --- | --- | --- | --- |
| DA59062 | 1-1 | 430 | *hisA*[Q18R A22T R23L K107E V168A] | WGS |
| DA59064 | 1-3 | 430 | *hisA*[Q18R A22T V168A] | WGS |
| DA59068 | 1-7 | 430 | (not identified)<sup>e</sup> | WGS |
| DA65429 | 2mut+3 | 78 | *trpA*[P62R(fs)] | WGS |
| DA65433 | 2mut+7 | 78 | *trpA*[P62R(fs)] | WGS |
| DA65434 | 2mut+8 | 78 | *trpA*[P62R(fs)] | WGS |
| DA65435 | 2mut+9 | 78 | *trpA*[P62R(fs)] | WGS |
| DA65436 | 2mut+10 | 78 | *trpA*[P62R(fs)] | WGS |
| DA65437 | 2mut+11 | 78 | *trpA*[P62R(fs)] | WGS |
| DA65438 | 2mut+12 | 78 | *trpA*[P62R(fs)] | WGS |
| DA65445 | 2mut+19 | 78 | *trpA*[P62R(fs)] | WGS |
| DA65447 | 2mut+21 | 78 | *trpA*[P62R(fs)], *trpA*[A67T] | WGS |
| DA65449 | 2mut+23 | 78 | *trpA*[P62R(fs)] | WGS |
| DA65451 | 2mutS1 | 78 | *hisA*[Q18R] | Sanger |
| DA65452 | 2mutS2 | 78 | *hisA*[Q18R A127G] | WGS |
| DA65453 | 2mutS3 | 78 | *hisA*[Q18R H17R] | WGS |
| DA65454 | 2mutS4 | 78 | *trpA*[D27Y G98S] | Sanger |
| DA65456 | 2mutS6 | 78 | *trpA*[P62R(fs)] | Sanger |
| DA65457 | 2mutS7 | 78 | *trpA*[P62R(fs)] | Sanger |
| DA65458 | 2mutS8 | 78 | *hisA*[Q18R A22T] (8-fold amplified)<sup>f</sup> | WGS |
| DA65459 | 2mutS9 | 78 | *hisA*[Q18R] | Sanger |
| DA65460 | 2mutS10 | 78 | *hisA*[Q18R] | Sanger |
| DA65461 | 2mutS11 | 78 | *trpA*[G98C] | Sanger |
| DA65462 | 2mutS12 | 78 | *trpA*[P62R(fs)] | WGS |
| DA65463 | 2mutS13 | 78 | *trpA*[P62R(fs)] | Sanger |
| DA65464 | 2mutS14 | 78 | *trpA*[G98S Y102H Q250R] | Sanger |
| DA65465 | 2mutS15 | 78 | *trpA*[P62R(fs)] | Sanger |
| DA65466 | 2mutS16 | 78 | *trpA*[T24S G61S P62S] | WGS |
| DA65467 | 2mutS17 | 78 | *hisA*[Q18R] | Sanger |
| DA65468 | 2mutS18 | 78 | *hisA*[Q18R] | WGS |
| DA65469 | 2mutS19 | 78 | *hisA*[Q18R T52S] | Sanger |
| DA65471 | 2mutS21 | 78 | *trpA*[P62R(fs)] | Sanger |
| DA65472 | 2mutS22 | 78 | (not identified)<sup>e</sup> | WGS |
| DA65473 | 2mutS23 | 78 | *trpA*[D27Y G98C] | WGS |

<sup>a</sup> Only populations that could grow significantly denser than their ancestor within 7 days at the end of the experiment are listed. The populations are named according to which evolution experiment (1 or 2) they were part of, and the order the cultures were handled in during passages (1 – 8 and 1 – 24, respectively for the first and second evolution experiment). The populations from the second experiment additionally have “mut+” for non-mutators and “mutS” for mutators.
<sup>b</sup> The numbers of generations were calculated from the fold dilution and the number of passages [ $\log_2(\text{dilution}) \times \text{passages}$ ] and reflect the minimum number of generations for the population as a whole.
<sup>c</sup> The mutations listed have been re-constructed in the genetic background of the non-evolved ancestral strain ( $\Delta trpF$ ), and has been verified to confer growth in the absence of tryptophan.
<sup>d</sup> Mutations were identified either through whole genome sequencing (WGS) or Sanger sequencing (targeting the *trpA* and *hisA* genes). For a complete list of mutations present at high frequencies in the whole-genome sequenced populations, see Table S1.
<sup>e</sup> These populations did not have any mutations in *hisA* or *trpA*.
<sup>f</sup> This population contained a ~19 kb amplification encompassing the entire *his* operon at ~8-fold coverage compared to the surrounding genomic DNA.

**Figure 2.**
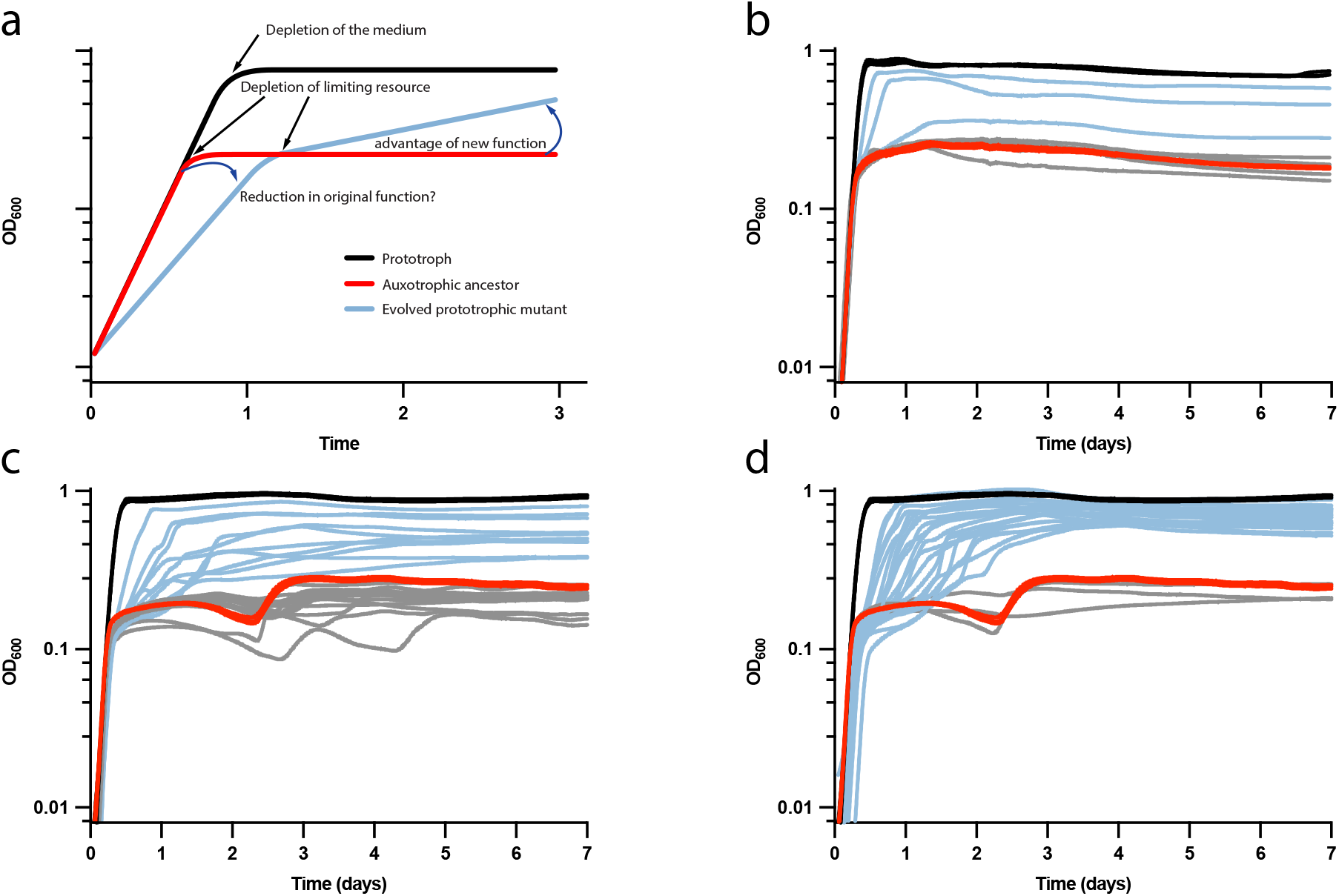
Growth Dynamics of Prototrophic and Auxotrophic Strains under Limiting Resources. **(a)** Hypothetical growth curves: the expected growth behaviour of a prototrophic strain (black), an auxotrophic strain (red), and an evolving prototroph (blue) under limiting resource conditions. The auxotroph (red) is expected to grow until the limiting resource is depleted and enter stationary phase at low population density. The prototroph (black) can continue growing until depletion of other essential nutrients or changes in medium composition (e.g., pH or metabolite accumulation) halt growth. The evolved prototroph (blue) exhibits growth after the limiting resource depletion, with a rate dependent on the rate of its newly acquired activity. If the original enzyme function is crucial for growth, reduced activity due to a mutation might lead to a slower growth rate before the limiting resource is depleted. **(b – d)** Batch growth curves of a tryptophan prototroph (*trpF*+, black), the ancestral tryptophan auxotroph (Δ*trpF*, red), the evolving populations from the final time-point that gained tryptophan independent growth (blue), and those that did not (grey). **(b)** The eight lineages from the first experiment. **(c)** The 24 non-mutator lineages from the second experiment. **(d)** The 24 mutator lineages from the second experiment (compared to the ancestral non-mutator strain).

The third population did not carry mutations in any readily identifiable target gene, and the genetic basis of its ability to grow after tryptophan depletion remains unresolved (Supplementary Results and Discussion: *Two populations evolved TrpF-independent growth in still unidentified ways*; Table S1).

### Mutator status influences both mutation frequency and genetic targets

To further assess the role of elevated mutation rates and to explore alternative evolutionary trajectories, a second experiment was performed in which 24 populations carrying a *mutS* mutation (mutators) and 24 populations without such a mutation (non-mutators) were evolved under the same intermittent selection regime for approximately 78 population doublings.

After this period, 21 mutator populations and 10 non-mutator populations exhibited detectable growth after tryptophan depletion (Figure 3c,d). WGS of a subset of populations, followed by targeted Sanger sequencing of the remaining populations, identified candidate mutations sufficient to confer the growth-rescue phenotype. These mutations were found either in *hisA* (nine mutator populations) or in *trpA* (11 mutator populations and 10 non-mutator populations).

**Figure 3.**
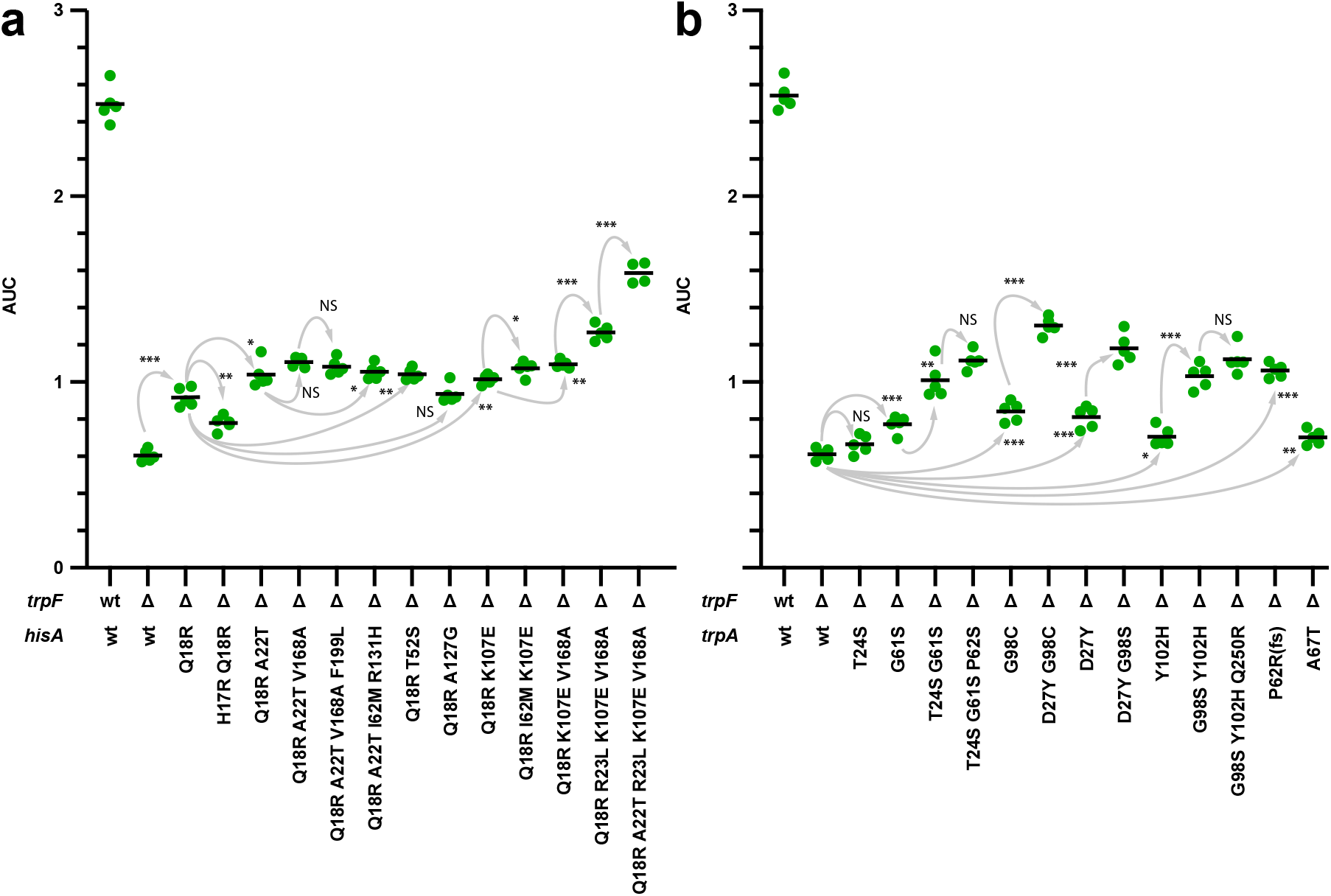
Growth of re-constructed mutants in tryptophan-limited medium. **(a)** *hisA* mutants. **(b)** *trpA* mutants. Reconstructed mutants were grown in M9 glucose minimal medium supplemented with limiting tryptophan (5 μM) and guanosine (3 mM). As growth under these conditions deviates from exponential growth (except for the TrpF+ control strain), growth is reported as the area under the curve (AUC) for the first 3 days (see Methods for details). Black bars represent averages. NS, non-significant (*p* > 0.05); *, *p* < 0.05; **, *p* < 0.005; ***, *p* < 0.0005, based on a two-tailed Student’s t-test with equal variance. Only the two rightmost *trpA* mutants were found in non-mutator populations, all others (in both genes) were found in hypermutators. Each mutant was compared to its immediate ancestor, indicated by the grey arrows. All strains were grown in at least four independent biological replicates. The complete growth curves can be found in Figures S1, S2, S3.

TrpA from *S. enterica* has previously been shown to support growth of Δ*trpF* strains through directed evolution and rational mutagenesis (Evran et al. 2012), but had not been identified as a target in prior non-directed evolution experiments. Notably, all sequenced non-mutator populations and six mutator populations shared the same *trpA* mutation: a 7-bp duplication causing a frameshift. For this mutation to be beneficial under the conditions used here, it must give rise to sufficient amounts of functional protein via ribosomal frameshifting that both substitutes for TrpF in vivo and retains sufficient TrpA function to support growth (see Supplementary Results and Discussion: *How can a frameshift mutation in trpA both create a new function and retain the original function?*). One non-mutator population additionally carried a second *trpA* mutation (A67T) in a subpopulation lacking the frameshift mutation. One mutator population did not acquire mutations in either *hisA* or *trpA*, and its mechanism of adaptation remains unexplained (Supplementary Results and Discussion: *Two populations evolved TrpF-independent growth in still unidentified ways*; Table S1).

Together, these results show that rescue of growth in Δ*trpF* strains can occur through mutations in at least two distinct genes, *hisA* and *trpA*, and that mutator mutations are not required for this outcome, although they influence the types and distributions of mutations observed.

### Gene duplication is rare and does not lead to detectable functional divergence

In the minimal medium used for the evolution experiments, growth after tryptophan depletion requires that both histidine and tryptophan biosynthesis remain functional. HisA function is required throughout the growth cycle, whereas TrpA function becomes necessary only after depletion of tryptophan. Despite these constraints, only one whole-genome– sequenced population harbored a duplication or amplification of a relevant target gene (*hisA*). This population carried a single *hisA* allele (Q18R A22T; Table S1) that was supported by 100% of reads mapping to the locus, with no evidence for additional divergent *hisA* alleles. Thus, although amplification occurred in this population, there was no detectable evidence that gene copies had begun to diverge into distinct functional roles.

### Evolutionary paths involving *hisA* and *trpA* differ in mutational structure

To determine the order of mutation accumulation in populations carrying multiple mutations in *hisA* or *trpA*, clones were isolated from earlier time points and the target genes were sequenced (Table S2). Consistent with previous work showing that single amino acid substitutions in HisA rarely enable growth of Δ*trpF* strains without compromising growth under histidine-limited conditions (Lundin et al. 2020), all evolutionary paths involving *hisA* began with the same mutation (Q18R).

In contrast, *trpA* exhibited greater diversity in initial mutations, suggesting a broader range of mutational paths that can support growth after tryptophan depletion. To isolate the effects of individual mutations from background mutations accumulated during evolution, all identified *hisA* and *trpA* variants from final populations, as well as most intermediate variants, were reconstructed in an otherwise unevolved genetic background. Growth after tryptophan depletion in these reconstructed strains therefore indicates that the evolved alleles were sufficient to confer the growth-rescue phenotype.

### Successive mutations improve growth under selection conditions

Growth of reconstructed strains was assessed in the same minimal medium used during the evolution experiments (Figures 3, S1, S2, S3). During these assays, an unexpected phenotype was observed in which *hisA* mutants grew more slowly in the presence of added tryptophan than in its absence unless guanosine was supplied (Supplementary Results and Discussion: *Supplementing the growth medium with tryptophan causes growth defects that can be overridden by guanosine*). This effect likely introduced additional selection pressures unrelated to rescue of tryptophan prototrophy during evolution.

To assess the effects of *hisA* mutations independent of this phenotype, guanosine was included in the growth medium. Under these conditions, growth after tryptophan depletion depends on maintenance of native HisA and TrpA functions together with the newly acquired ability to substitute for TrpF *in vivo*. All reconstructed *hisA* and *trpA* variants enabled growth after tryptophan depletion (Figures S1–S3). In most reconstructed mutational pathways, each additional mutation resulted in improved growth following tryptophan depletion.

### Trade-offs with ancestral function differ between *hisA* and *trpA* variants

Prior to tryptophan depletion, all reconstructed *trpA* mutants grew at rates indistinguishable from the ancestral strain, which itself grew similarly to a TrpF^+^ prototrophic strain (Figures S3 and S4). This is expected because TrpA function is not required when tryptophan is present in the medium.

In contrast, all reconstructed *hisA* mutants displayed reduced growth rates relative to the ancestral strain, and successive *hisA* mutations further decreased growth (Figures 4a, S1, S2). Although supplementation with guanosine partially restored growth, all *hisA* mutants still grew significantly more slowly than a wild-type *hisA* control (Figure 4a), consistent with a decline in the native HisA function. As shown previously (Newton et al. 2017), this decline likely reflects combined effects on catalytic properties and other factors that influence total enzyme output *in vivo*.

**Figure 4.**
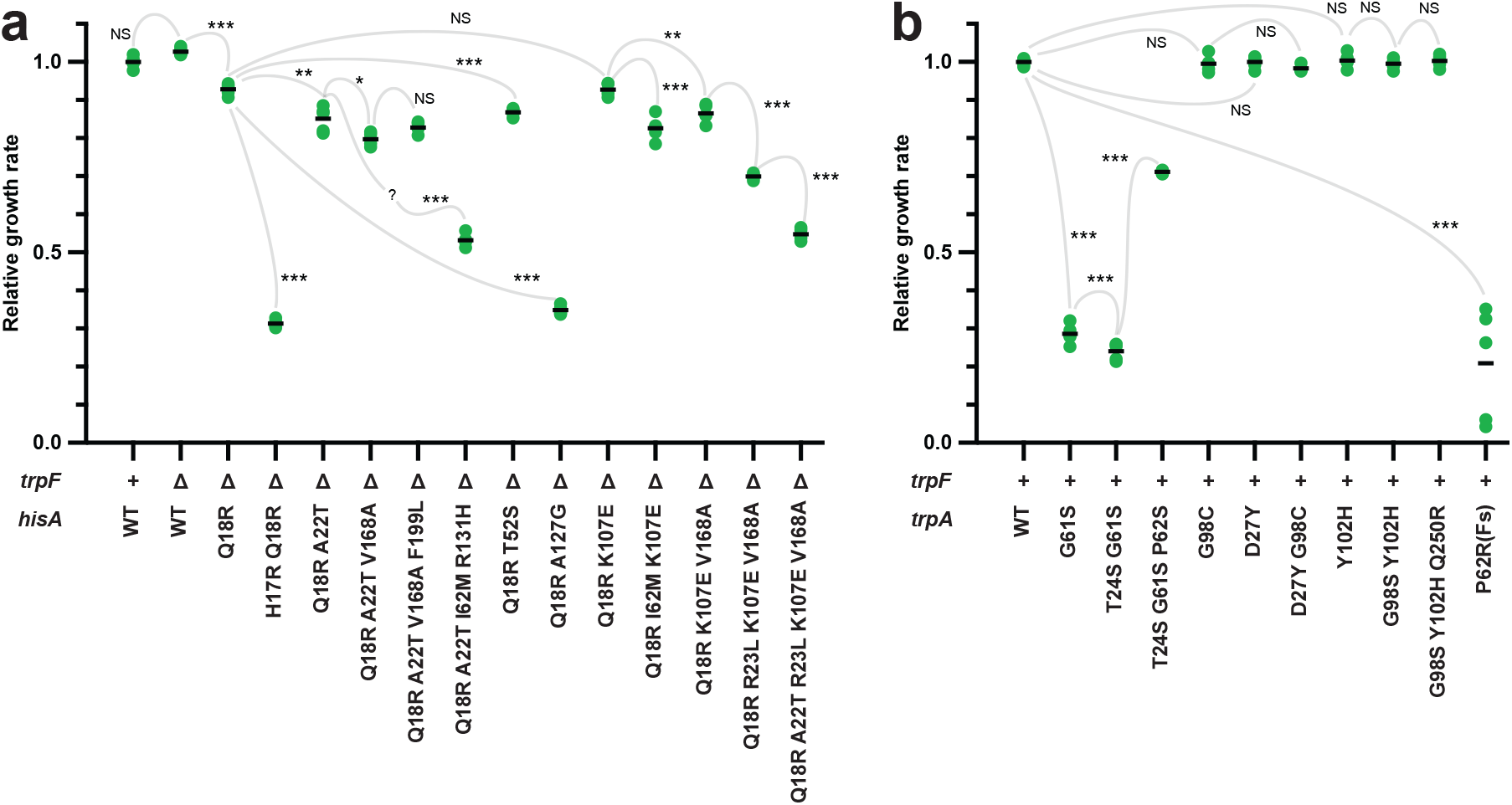
Impact of Mutations on native function. **(a)** HisA function: Exponential growth rates were measured in the early, rapid growth phase (0.095 < OD_600_ < 0.184) before tryptophan depletion in M9 glucose medium supplemented with limiting tryptophan (5 μM) and guanosine (3 mM). Rates were normalized to the growth rate of the *trpF*+ wild-type strain grown in the same experiment. Under these conditions, reductions in HisA activity lead to decreased growth rates. Strains were grown in four or five biological replicates. Grey lines connect each mutant to its immediate ancestor. The question mark indicates an unknown mutation order in the evolution experiment, preventing complete reconstruction of the path. **(b)** TrpA function: Exponential growth rates were measured in M9 glucose medium without supplements for strains expressing wild-type *trpF* from a constitutive promoter at a different locus. Here, reductions in TrpA function lead to decreased growth rates. For *trpA* mutants under the same conditions as (a), see Figure S4, and for growth curves of strains complemented with *trpF* or *trpA*, see Figure S5. NS, non-significant (difference in growth rate < 5% or p > 0.05). Asterisks indicate significance levels (*, *p* < 0.05; **, *p* < 0.005; ***, *p* < 0.0005) based on two-tailed Student’s *t*-tests with equal variance, comparing each mutant to its immediate ancestor (grey lines).

Because TrpA and TrpF act within the same biosynthetic pathway, growth curves of *trpA* mutants alone cannot distinguish between changes in the two functions. To assess effects on native TrpA-dependent growth, a wild-type *trpF* gene was expressed from a strong constitutive promoter at an unrelated locus in ten reconstructed *trpA* mutants. Under these conditions, six mutants displayed growth indistinguishable from wild-type controls, indicating minimal impairment of TrpA-dependent growth (Figure 4b). Four mutants grew more slowly, consistent with reduced TrpA function.

To further evaluate TrpA-dependent growth capacity, wild-type *trpA* and *trpB* were co-expressed from the same locus. Expression of TrpAB improved growth after tryptophan depletion but did not restore growth to the same extent as expression of TrpF, indicating that the TrpF-substituting function was more limiting under these conditions (Figure S5). One growth-compromised variant carried the P62fs frameshift mutation, which is expected to yield low levels of functional protein due to reliance on frameshift suppression. Consistent with this, TrpAB complementation had little effect in this background.

Three *trpA* variants originated from the same evolutionary pathway in which the initial mutation (G61S) enabled growth of Δ*trpF* strains but impaired TrpA-dependent growth, a second mutation (T24S) improved growth after tryptophan depletion without further loss of TrpA function, and a third mutation (P62S) restored TrpA-dependent growth (Figure S5d–f).

### Summary of Results

Together, these results show that growth of Δ*trpF* strains under intermittent selection can be restored through mutations in multiple genes without gene duplication and divergence. Mutations in *hisA* and *trpA* both supported rescue of tryptophan prototrophy, but differed in their mutational accessibility and in the extent to which ancestral growth-supporting functions were compromised. These findings provide an empirical basis for understanding how gene-level multifunctionality can emerge during adaptation without gene duplication.

## Discussion

This study was designed to identify genetic solutions that restore growth of Δ*trpF* bacteria under conditions intended to minimize experimental bias. In contrast to earlier work that used engineered starting alleles, altered expression, amplification-prone loci, or targeted mutagenesis (Jürgens et al. 2000; Evran et al. 2012; Näsvall et al. 2012; Lundin et al. 2020), we initiated evolution from bacteria carrying wild-type alleles of *hisA, trpA*, and other relevant genes in their native genomic and regulatory contexts. Populations were propagated in medium containing a limiting amount of tryptophan, producing intermittent selection for rescue of tryptophan prototrophy as tryptophan became depleted each growth cycle. Under these conditions, mutations in either *hisA* or *trpA* repeatedly enabled growth after tryptophan depletion, and reconstruction of evolved alleles showed that these mutations were sufficient to confer the growth-rescue phenotype (Figure 3b–d; Tables 1, S1, S2; Figures S1–S3).

### Multiple genes provide genetic solutions to loss of *trpF* under minimally manipulated conditions

A central result is that Δ*trpF* growth rescue emerged via mutations in at least two different genes, *hisA* and *trpA*, across independent populations (Figure 3; Tables 1 and S1). Prior literature provides clear biochemical precedent that both enzymes can acquire TrpF catalytic activity measurable *in vitro* (Jürgens et al. 2000; Evran et al. 2012; Newton et al. 2017). In addition, previous work has shown that some HisA variants can be bifunctional at the protein level, retaining measurable native activity while also catalyzing the TrpF reaction *in vitro* (Jürgens et al. 2000; Newton et al. 2017), and natural HisA orthologs (PriA) can support both histidine and tryptophan biosynthesis *in vivo* (Barona-Gómez and Hodgson 2003; Due et al. 2011; Plach et al. 2016). Consistent with these earlier findings, the mutations identified here repeatedly generated *genes* that support growth in the absence of *trpF* while retaining sufficient ancestral function to support growth under the same conditions.

Importantly, our primary readout is physiological: the ability of reconstructed alleles to support growth after tryptophan depletion in Δ*trpF* strains. Throughout, “function” refers to this in vivo growth support rather than catalytic rate constants or enzyme kinetics. Accordingly, the central conclusions of this study do not depend on the exact biochemical mechanisms underlying growth rescue, but on the genetic and phenotypic observations that specific evolved *hisA* and *trpA* alleles are sufficient to restore growth in the absence of *trpF* while retaining enough ancestral growth-supporting capacity under the same conditions. This framing is aligned with the objectives of the study—mapping genetic changes to adaptive growth phenotypes—and avoids over-interpreting growth-based outcomes as direct measures of catalytic efficiency.

### Mutational accessibility and trade-offs differ between *hisA* and *trpA* solutions

Although both genes repeatedly yielded growth-rescuing alleles, the evolutionary paths differed. In Experiment 1, *hisA* mutations were found only in populations where mutator alleles appeared prior to the first *hisA* mutations (Tables S1, S2), and in Experiment 2, *hisA* solutions again occurred primarily in *mutS* backgrounds (Figure 2c, d; Table S1). This pattern suggests that, under the conditions used here, mutator status increased access to *hisA*-based solutions, either by elevating the overall supply of rare beneficial mutations or by enabling multi-step paths to be sampled on the relevant timescale. Consistent with this, all reconstructed *hisA* mutants showed reduced growth relative to wild-type *hisA* controls even when guanosine was supplied (Figure 4a), indicating a measurable decline in the ancestral growth-supporting function of HisA.

In contrast, *trpA* solutions were common in both mutator and non-mutator backgrounds (Figure 2c,d; Table S1). Several reconstructed *trpA* variants displayed wild-type growth rates when *trpF* was provided in trans (Figure 4b), consistent with little or no reduction in TrpA-function under those conditions. Other *trpA* variants showed reduced growth that could be improved by providing *trpAB* in trans (Figure S5), indicating that impairment of TrpA function contributed to the phenotype for at least some alleles. Together, these results support the view that *hisA*-based solutions more frequently incur detectable trade-offs with ancestral growth support than *trpA*-based solutions in this experimental context.

One striking outcome was the repeated appearance of a 7-bp duplication causing a frameshift in *trpA* in many non-mutator populations (and in several mutator populations) (Table S1). As discussed in the Supplementary Results and Discussion: *How can a frameshift mutation in trpA both create a new function and retain the original function?*, this allele must yield sufficient functional protein through ribosomal frameshifting to support growth. Regardless of the molecular basis, the recurrence of this mutation indicates that it is an accessible and competitive route to Δ*trpF* growth rescue under the conditions used here.

### Gene duplication was rare and did not lead to detectable divergence

A second notable difference from prior work is the low prevalence of gene duplication or amplification. Only one population showed amplification of a relevant target gene (*hisA*) (Table S1), and in this population the reads supported a single *hisA* allele without evidence of divergent gene copies. This contrasts with earlier experimental designs that used amplification-prone genomic contexts and constitutive expression, where duplication and subsequent divergence were central features (Näsvall et al. 2012). Several aspects of the present design could contribute to this difference, including native chromosomal context and regulation of the evolving genes, wild-type starting alleles rather than pre-adapted mutants, and a selective regime in which selection for growth rescue occurs only after tryptophan depletion in each cycle.

While we cannot exclude the possibility that transient duplications arose and were lost during evolution, the absence of duplications in the vast majority of end-point populations, coupled with the absence of evidence for divergent alleles in the single amplified population, indicates that duplication was not a dominant route to adaptation in these experiments. This is consistent with the broader expectation that many duplications carry fitness costs and are genetically unstable, requiring sustained positive selection to be maintained (Hill et al. 1969; Haack and Roth 1995; Reams et al. 2010). Under intermittent selection, any benefit of duplication that is restricted to the post-depletion phase could be reduced by loss during phases lacking selection for growth rescue, potentially favoring point-mutation routes that produce single genes capable of supporting both requirements.

### Intermittent selection creates a stringent and time-dependent filter on adaptive solutions

Growth under the evolution regime reflects more than the ability to resume growth after tryptophan depletion. Even slow-growing or non-growing cells continue to consume resources after depletion of tryptophan, making the time available for additional growth constrained by depletion of other limiting resources. Consequently, a growth-rescue mutation must enable sufficient post-depletion growth to compensate for any pre-depletion deficit and to outcompete other genotypes before the medium becomes exhausted. This time-dependent filter may limit the set of mutations that are not only beneficial in principle but also competitively successful under the specific cycle structure.

More generally, fluctuating selection is known to promote generalist phenotypes at the organismal level, particularly when adaptation involves trade-offs across environments (Kassen 2002; Legros and Koella 2010; Hall et al. 2013; Arribas et al. 2014; Kortright et al. 2022; Fasanello et al. 2024). The present results are consistent with the idea that a similar logic can apply at the gene level: when selection alternates between phases emphasizing different requirements, evolution may favor alleles that satisfy multiple requirements simultaneously, even if those alleles are not optimal for either requirement alone. The cycle structure and frequency of environmental switching are known to influence these outcomes (Roth and Maisnier-Patin 2016; Van Hofwegen et al. 2016), and in this system the alternation between growth with exogenous tryptophan and growth without it provides a clear, repeated switch in selection pressures.

### Caveats and interpretation of “bifunctionality” at the gene level

A key point of framing is that our conclusions are based on gene-level phenotypes. The reconstructed *hisA* and *trpA* alleles were sufficient to confer growth after tryptophan depletion in Δ*trpF* strains, while still permitting growth under conditions requiring the ancestral functions (Figures 4, S1–S3). This establishes bifunctionality at the level of *genes supporting two physiological requirements*. It does not, by itself, quantify catalytic activities or identify the biochemical mechanism.

That said, several lines of evidence support the interpretation that the most parsimonious explanation is acquisition of TrpF-like catalytic activity by the mutant proteins while retaining native catalytic function, consistent with prior literature (See Figure S7 for the reactions catalyzed by the three enzymes). First, HisA and TrpA variants that rescue Δ*trpF* in other studies have been shown to possess TrpF catalytic activity *in vitro* (Jürgens et al. 2000; Evran et al. 2012; Newton et al. 2017). Second, some HisA variants with TrpF catalytic activity retain measurable native activity *in vitro* (Jürgens et al. 2000; Newton et al. 2017). Third, in the present study, most *trpA* variants displayed wild-type growth when complemented with *trpF* (Figure 4b), supporting retention of TrpA function *in vivo* for many alleles. Conversely, where complementation experiments indicated partial deficits, they are consistent with altered TrpA function rather than bypass of the pathway (Figure S5). For a speculative discussion on potential mechanisms of action of the evolved HisA and TrpA proteins, see Supplementary Results and Discussion: *Speculative considerations for ΔtrpF rescue by evolved HisA and TrpA proteins* and Figure S8. While alternative mechanisms cannot be formally excluded, the combination of genetic sufficiency and the strong precedent for enzymatic acquisition in this system supports the interpretation that the mutant proteins are capable of catalyzing both their native reactions and that they can substitute for the missing TrpF.

## Conclusions

By evolving Δ*trpF* populations under a selective regime designed to reduce experimental bias, we found that point mutations in multiple genes (*hisA* and *trpA*) can repeatedly rescue growth without exogenous tryptophan while retaining sufficient ancestral growth-supporting functions. Mutational paths and trade-offs differed between genes, and gene duplication was rare and showed no evidence of divergence. Together with prior biochemical and comparative evidence (Jürgens et al. 2000; Barona-Gómez and Hodgson 2003; Due et al. 2011; Evran et al. 2012; Plach et al. 2016; Newton et al. 2017), these results support the broader conclusion that gene-level multifunctionality can emerge during adaptation without gene duplication, and that the structure of selection—here, intermittent selection imposed by nutrient depletion—can shape which genetic solutions are accessible and successful.

## Data availability

All genome re-sequencing data have been deposited in the Sequence Read Archive (SRA) with BioProject accession code PRJNA1092712.

## Acknowledgements

We thank Anna Knöppel, Omar Warsi, Ramith Nair, and Dan Andersson for critical reading of the manuscript.

## Author Contributions

JN conceived the study, constructed bacterial strains, made bacterial growth rate assays and analyzed data. HA and JN performed evolution experiments, and isolated and sequenced clones from evolved populations. HA wrote an early draft of the manuscript as part of her PhD thesis, JN re-worked and extended the manuscript.

## Supplementary Information

Supplementary Information is available for this paper.

