## Supplemental Text and Figures for "Experimental evolution reveals bifunctional genetic solutions to loss of *trpF* in *Salmonella enterica*"

Supplementary Materials and Methods
Supplementary Results and Discussion
Supplementary figures S1 – S8

### Supplementary Materials and Methods

#### Handling of phage P22 and *gal* operon modifications

Generalized transduction between *Salmonella enterica* strains was primarily performed using phage P22 *HT105/1 int-201* (Schmieger 1972). In long-term evolution experiments and repeated transductions, contamination by P22 or selection for P22-resistant mutants can complicate downstream genetic analyses. To minimize these issues, two complementary strategies were used.

In evolution experiment 1, strains carried deletions in the *gal* operon (Δ$P_{gal}$ Δ*galE*::$T_{lux}$). GalE catalyzes the reversible interconversion of UDP-galactose and UDP-glucose, and loss of GalE results in conditional truncation of the O-antigen in the absence of galactose. Because phage P22 uses the O-antigen as a receptor, this modification confers resistance to P22 infection in the absence of galactose while maintaining sensitivity in its presence. To avoid galactose sensitivity due to toxic accumulation of UDP-galactose, the *gal* operon promoter was deleted and a transcriptional terminator was inserted in place of *galE*, reducing expression of *galK* and *galT*. The resulting strain was resistant to P22 in the absence of galactose, sensitive in its presence, and showed no detectable galactose sensitivity. These *gal* mutations were used to reduce the likelihood of P22 contamination and to avoid selection for P22-resistant mutants during the evolution experiments.

#### Construction and use of an artificial Gene Transfer Agent (GTA22)

To facilitate repeated transductions while preventing lytic growth of phage P22, an inducible artificial gene transfer agent (GTA), here referred to as GTA22, was constructed using λ Red recombineering. GTA22 was derived from a P22 *HT105/1 sieA44* lysogen by three major genetic modifications.

First, both ends of the prophage, including the attachment sites, were deleted using DIRex (Näsvall 2017; Näsvall 2022). One deletion (Δ*thrW*–*kil*) removed approximately 6.9 kb from the left end of the prophage, including the left attachment site within the *thrW* tRNA gene, the integrase and excisionase genes (*int* and *xis*), recombination genes (*abc1*, *abc2*, *erf*), and the septation inhibitor gene *kil*. A second deletion removed approximately 3 kb from the right end of the prophage, including the right attachment site (a partial duplicate of *thrW*) and the O-antigen conversion locus *gtrABC*. The *gtrABC* locus mediates phase-variable modification of the O-antigen, which can prevent adsorption of P22 and thereby block transduction.

Second, to make the prophage inducible for production of transducing particles, a copy of the P22 antirepressor gene ($ant_{P22}$) was placed under control of the arabinose-inducible $P_{araBAD}$ promoter in the host chromosome using λ Red recombineering. In the absence of attachment sites and excision functions, induction of $ant_{P22}$ allows replication from the phage origin to extend into adjacent host DNA without excision or circularization of the prophage genome.

Because packaging of P22 DNA into virions initiates at a single *pac* site and proceeds unidirectionally (Susskind and Botstein 1978), GTA22 cannot package a complete copy of its own genome and is therefore unable to form viable phage particles under normal conditions. Although rare formation of virulent particles via illegitimate recombination cannot be formally excluded, GTA22 behaved as avirulent in all experiments performed.

Strains carrying GTA22 retained the P22 repressor (c2) and were resistant to lytic infection by superinfecting P22 genomes while remaining competent both as donors and recipients in generalized transductions. This eliminated the need for extensive screening to exclude phage-contaminated clones.

To induce GTA22, overnight cultures were diluted 1:20 into LB supplemented with 0.1% (w/v) L-arabinose and incubated for at least 6 h at 37 °C to allow lysis. Surviving bacteria were killed by vigorous shaking with chloroform. Lysates were clarified by centrifugation (≥14,000 × g for 1–3 min) and used directly in transductions. For transduction, 0.1–10 µL of cleared lysate was mixed with 100 µL LB supplemented with 0.4% glucose to repress $P_{araBAD}$ in recipient cells, followed by addition of 100 µL overnight culture of the recipient strain. After 30 min incubation at 37 °C, mixtures were plated on selective media.

#### Isolation and sequencing of clones from evolved populations

To determine the order of appearance of mutations in populations that accumulated multiple mutations in *hisA* or *trpA*, clones were isolated from frozen population samples taken at earlier time points. Frozen stocks were streaked on LA plates, and eight single colonies per population were restreaked once more for purification. From each purified clone, a single colony was used as template for PCR amplification of the target gene, followed by Sanger sequencing. Selected clones were archived in the strain collection and used as templates for reconstruction of individual mutations in unevolved genetic backgrounds.

### Additional details of growth assays and data handling

During Bioscreen growth assays, evaporation of medium from wells at the edges of the plates was occasionally observed despite sealing of the plates. In early experiments, edge wells sometimes reached higher final OD values and showed visible volume loss after prolonged incubation. This effect became more pronounced at later time points and contributed to increased variation among replicates. However, up to approximately 90% of maximum OD (prior to entry into stationary phase), evaporation did not appear to substantially affect growth measurements, as affected and unaffected wells showed similar OD values up to this point.

In later experiments, edge wells were filled with medium but not used for cultures to minimize evaporation effects. For fitness comparisons based on area under the curve (AUC), analyses were restricted to the first 72 h of growth, which avoided most artefactual variation due to evaporation.

Some strains, particularly the ancestral strain of evolution experiment 2 and reconstructed derivatives, displayed fluctuations in OD600 following depletion of tryptophan. After an initial period without growth, OD600 sometimes decreased before increasing again. These fluctuations occurred within the first three days and are interpreted as reflecting entry into death phase followed by partial regrowth due to recycling of nutrients from lysed cells. Because this behaviour appeared independent of post-depletion growth rescue, it likely inflates AUC values for strains with little or no growth rescue, thereby underestimating differences between strains with and without growth rescue activity.

### Generation of a mutator allele

A reversible mutator allele was constructed by insertion of the selectable and counter-selectable *Acatsac1* cassette (GenBank accession MF124798) into *mutS* using a duplication–insertion (DUP-In) strategy (Näsvall et al. 2017). The insertion generated two truncated copies of *mutS*, each lacking essential parts of the *mutS* coding sequence, separated by the *Acatsac1* cassette. As neither truncated copy is expected to encode a functional MutS protein, the resulting strains displayed a mutator phenotype, as confirmed by elevated mutation counts in evolved populations. The DUP-In design allows reversion of the mutator allele by recombination between duplicated sequences and loss of the cassette upon selection on sucrose, although reversion was not tested in this study.

### Constitutive expression of *trpA* and *trpF* from a heterologous locus

To assess the effects of *trpA* mutations on TrpA and TrpF functions separately, wild-type *trpA* and *trpF* genes were expressed from the strong constitutive $P_{CP25}$ promoter (Jensen and Hammer 1998) at a chromosomal locus unrelated to the native tryptophan biosynthetic operon. Recipient strains carried a $P_{CP25}$-driven Red fluorescent protein (mScarlet) gene inserted at one of the six IS*200* elements in the *S. enterica* genome, along with deletions of either *trpA* or *trpF*. λ Red recombineering was used to replace the mScarlet coding sequence with *trpA* or *trpF*, selecting for growth in the absence of tryptophan.

To facilitate transfer of the $P_{CP25}$-*trpA* and $P_{CP25}$-*trpF* constructs to other strains, a duplication containing a selectable and counter-selectable marker (*dhfr*–$P_{rhaB}$–*orph11*) was inserted adjacent to the same locus. Following generalized transduction and selection for trimethoprim resistance, loss of the marker was selected by growth in the presence of rhamnose, yielding marker-free strains.

Initial tests showed that $P_{CP25}$-*trpA* fully complemented a Δ*trpA* (*trpF*$^{+}$) strain but failed to restore growth of reconstructed *trpA* (Δ*trpF*) mutants. Because TrpA functions in concert with TrpB as an $\alpha_2\beta_2$ complex (Pittard and Yang 2008), competition between wild-type and mutant TrpA proteins for TrpB binding may inhibit the function of the mutant TrpA variants. To address this, a $P_{CP25}$-*trpBA* construct co-expressing *trpB* and *trpA* was generated. This construct fully complemented Δ*trpA* (*trpF*$^{+}$) strains and partially complemented *trpA* (Δ*trpF*) mutants with reduced TrpA function, consistent with restoration of TrpA function while growth remained limited by the extent of Δ*trpF* growth rescue (Figure S6).

### Copy number estimation and correction of allele-frequency estimation from read depth

Copy number in the population containing amplification of a relevant target gene (*hisA*) was estimated from mapped read depth by dividing the mean read depth across structural genes in the *his* operon (7136 bp; 156.49×) by the mean read depth across four identically sized flanking regions (4 × 7136 bp; mean 19.95×).

For the *trpA*(Pro62Fs) mutation, CLC Genomics Workbench initially reported the variant in only 30–60% of reads at the locus. Visual inspection of the mapped reads revealed that a fraction of reads containing the mutation (a 7 bp tandem duplication) were classified as broken-ended reads rather than mutant reads, leading to underestimation of the true mutant allele frequency at this position. To correct for this, reads were re-mapped to a reference sequence carrying the frameshift allele; under this mapping, 82–100% of reads from the relevant populations mapped perfectly to the mutation.

### Supplementary Results and Discussion

#### Mutators precede the first *hisA* mutations

Evolution experiment 1 ran for approximately 430 population doublings, while experiment 2 ran for only 78 doublings. To compare the outcomes at similar time scales, we sequenced the original populations (1-1 through 1-8) from cycle 15 (approx. 100 generations; Table S1).

As expected, the populations had fewer mutations at this early stage, and none of the *hisA* or *trpF* mutations were detected. Most populations showed mutations that were present at the end-point, while other mutations that were detected early were not detected later.

These results indicate that the populations had already begun evolving and diverging into multiple interfering clones early on. However, no mutations conferring post-depletion growth had emerged or accumulated to detectable frequencies.

Interestingly, the mutator alleles (*mutH* and *mutS*) that were fixed in the end-points of populations 1-1 and 1-3, respectively, were present at low frequencies early on. This suggests that the mutators arose before the *hisA*(Q18R) mutation, which was detected by Sanger sequencing five cycles (about 33 generations) later (Table S2). These results reinforce our notion that mutators were essential for the evolution of restored prototrophy through single nucleotide substitutions under these conditions.

#### Supplementing the growth medium with tryptophan causes growth defects that can be overridden by guanosine

The HisA variants with the most Δ*trpF* rescue activity displayed faster growth in the absence of tryptophan compared to its presence (Figures S2 and S3). This observation might be linked to the availability of phosphoribosylpyrophosphate (PRPP), a precursor molecule common to the *de novo* synthesis of histidine, tryptophan, purines, pyrimidines, and pyridine nucleotides.

In wild-type *Salmonella enterica*, histidine feedback-inhibits the first enzyme (HisG) of its biosynthetic pathway, making HisG the committing and rate-determining step (Winkler and Ramos-Montañez 2009). Conversely, a mutation in any other enzyme besides HisG is expected to decrease histidine synthesis, leading to increased flux into the pathway due to derepression of the operon and reduced HisG inhibition. Consequently, a HisA mutation could potentially increase PRPP consumption for histidine biosynthesis, limiting its availability for other pathways.

Tryptophan addition slightly reduces the PRPP pool in wild-type *S. enterica*, possibly through indirect effects on other pathways (Sadler and Switzer 1977). A similar phenomenon was observed in *S. enterica* mutants with a transposon insertion in *trpC*, exhibiting a filamentous growth phenotype. This phenotype was reversed by the addition of histidine, tryptophan, mutations in *hisF* or *hisH*, but not by nucleotides (Henry et al. 2005). This effect was attributed to excessive PRPP channeling into tryptophan synthesis, leaving insufficient PRPP for histidine biosynthesis.

We hypothesized that the observed growth inhibition of HisA mutants upon tryptophan addition might be related to its effect on the PRPP pool. Reduced PRPP synthesis due to tryptophan addition, coupled with increased PRPP channeling into the histidine pathway due to the HisA mutation, could potentially lead to insufficient nucleotide synthesis.

Given the critical role of purines (adenosine and guanosine) in energy homeostasis and as building blocks for RNA and DNA, compared to the less demanding pyrimidines (cytosine, uridine, and thymidine(Buckstein et al. 2008), we reasoned that if the growth reduction stemmed from limited PRPP availability, supplementing the medium with guanosine could rescue the growth defect. Notably, guanosine addition improved the growth of HisA mutants in tryptophan-containing medium but had no significant effect on wild-type HisA strains (including *trpA* mutants).

Regardless of the precise mechanism underlying the growth reduction in HisA mutants upon tryptophan addition, it appears unrelated to a direct limitation in histidine or tryptophan biosynthesis. Therefore, to isolate the specific effects on the native function of *hisA*, guanosine was added to the growth media when assaying the growth of reconstructed mutants in the presence of tryptophan (Figures 3, 4, S3, S4, S5, S6).

### Two populations evolved TrpF-independent growth in still unidentified ways

Two populations (designated "1-7" and "2*mutS*22" in Table 1) evolved the ability for TrpF-independent growth through unidentified mechanisms.

Identification of the specific mutations responsible for this phenotype in these populations proved challenging. While both populations shared four mutation targets (in *yafS*, *yeaG*, *trpD*, and large duplications between rRNA operons), these alterations were also present in other populations, making them unlikely candidates for TrpF-independent growth.

The involvement of *trpD* mutations in the phenotype was specifically excluded. We reintroduced the wild-type allele of *trpD* into a clone isolated from population "2*mutS*22" using transduction. Despite this genetic modification, the transductants retained their ability for TrpF-independent growth, regardless of the *trpD* allele they possessed.

Furthermore, although isolated clones from population "2*mutS*22" functioned effectively as both a donors and recipients in transduction experiments, the TrpF-independent growth phenotype could not be transferred to the ancestral strain through this method. This suggests a polygenic basis for the phenotype, where mutations in multiple genes contribute, each with an effect too subtle for successful selection during single-gene transduction.

### How can a frameshift mutation in *trpA* both create a new function and retain the original function?

The isolated frameshift mutation in *trpA* (P62fs; dup173 – 179; Figure S6) is a seven-nucleotide duplication that disrupts the reading frame after codon 60. This leads to the misincorporation of 23 incorrect amino acids before translation terminates at a UGA stop codon. Given the early location of the mutation, the resulting 83-amino acid peptide is highly unlikely to retain any functional activity.

For the P62fs mutant to acquire a new function, the frameshift mutation in *trpA* requires efficient frameshift suppression. This suppression would allow translation to resume in a new reading frame, generating a full-length protein with an altered amino acid sequence.

Intriguingly, several populations harboring the P62fs mutation in *trpA* also displayed mutations in genes encoding ribosomal components: rRNA methyltransferase RsmD (YhhF) and ribosomal proteins S2 and L9 (Table S1). Mutations in L9 are known to enhance ribosomal frameshifting (Leipuviene and Björk 2007). While a direct role for RsmD in frameshifting has not been established, it contributes to frameshift suppression when combined with another frameshift suppressor mutation (Arora et al. 2013). Notably, both populations with RsmD mutations also possess mutations in S2. This co-occurrence suggests a potential synergistic effect of S2 and RsmD mutations in suppressing the *trpA* frameshift mutation.

### Speculative considerations for Δ*trpF* rescue by evolved HisA and TrpA proteins

This section provides speculative structural interpretations that are consistent with the observed ability of evolved *hisA* and *trpA* alleles to support growth of Δ*trpF* strains. These considerations are not required for the conclusions of this study and are not intended as evidence for specific catalytic mechanisms.

The enzymes TrpF, HisA, and TrpA are structurally related $(\beta\alpha)_8$-barrel proteins that act on similar substrates. Both TrpF and HisA catalyze Amadori rearrangements on their amino aldose substrates to form amino ketoses (Figure S7). Their catalytic mechanisms involve protonation of the ring oxygen of the reacting ribose by a general acid (D176 in HisA, D126 in TrpF) and deprotonation of the ribose 2' carbon by a general base (D7 in HisA, C7 in TrpF; Figure S7d).

For the TrpA reaction, a base (D60) abstracts a proton from the indole ring nitrogen (N1), while another residue (E49) acts as both catalytic acid and base that protonates the C3 position of the indole ring and deprotonates the hydroxyl of the leaving group (Figure S7e). Both HisA and TrpA have flexible loops that close over the active site after substrate binding (Söderholm et al. 2015; Duran et al. 2024 Nov 2). In TrpA, loop dynamics are critical for allosteric regulation by TrpB (Duran et al. 2024 Nov 2). In HisA, loop closure coincides with the rearrangement of the substrate into an extended product-like conformation (Söderholm et al. 2015).

Evran *et al.* (Evran et al. 2012) employed rational design to introduce putative catalytic residues from TrpF into TrpA, but only achieved TrpF activity after extensive mutagenesis and gene shuffling. Notably, their proposed essential mutations (F22C and L177D) were not sufficient for detectable TrpF activity. In contrast, our evolutionary approach identified several single amino acid substitutions (G98C, D27Y, G61S, Y102H) that were sufficient for rescue of Δ*trpF*, without introducing residues previously proposed to be essential for TrpF catalysis by mutant TrpA. Interestingly, some mutations identified by Evran *et al.* (Evran et al. 2012) as "permissive" (G61S, T24S) were sufficient for Δ*trpF* growth rescue in our experiments, suggesting these mutations may play a more significant role than previously thought, and that *trpA* mutations can support Δ*trpF* growth without mutations that exactly transplant the TrpF active site into TrpA.

Superposing the crystal structures of TrpA with its native substrate and TrpF with its product analog rCdRP places the carboxyl group of TrpA E49 and the sulfide group of TrpF C7 close to each other. Interestingly, TrpA E49 is only 3Å from the C2' of rCdRP, placing it in a position that could, in principle, participate in chemistry analogous to that required for the TrpF reaction. The respective general acids are far from each other, and TrpA D60 does not appear to be appropriately positioned for acting as general acid on PRA (Figure S8f). The absence of appropriately located potential general acids for the TrpF reaction in TrpA in the superposed structures may not necessarily be a relevant problem; in the structure of the bifunctional HisA ortholog PriA from *Mycobacterium tuberculosis* the product CdRP binds in a different orientation than in the TrpF structure, more resembling the binding of ProFAR in the active site of HisA (Figures S8d and S8e, (Due et al. 2011). This observation suggests the possibility that if PRA binds to the active site of TrpA, it does not necessarily bind in an orientation similar to its binding in TrpF (or PriA), but rather that it needs to bind in a conformation similar to the native TrpA substrate.

Several TrpA mutations that resulted in Δ*trpF* growth rescue map to positions close to the active site and could potentially affect the local structure or dynamics around the catalytic residues D60 or E49: G61S and P62S affect residues next to D60; Y102, close to D60, likely helps lock it in place for native catalysis, and the Y102H mutation may allow more flexibility or make the active site cavity larger; T24, D27, and A67 pack against each other at the interface between loop 1 and a helix at the end of loop 2. Mutations in these residues (T24S, D27Y, and A67T) may disrupt this interaction, leading to more flexibility of loop 2, perhaps allowing D60 to reach the ribose ring oxygen of PRA. G98 is close to E49, below the substrate binding pocket. Mutations at this position (G98C and G98S) add bulky side-chains that may restructure the active site to allow binding or catalysis of the larger TrpF substrate. One mutation (Q250R) maps to the outside surface far from the active site, making any speculation about its effect difficult.

In a previous evolution experiment testing the Innovation – Amplification – Divergence model for the evolution of new functions (Näsvall et al. 2012), we found two mutations in HisA that generated TrpF activity at the expense of all detectable HisA activity: "dup13-15" and L169R. The former results in a three-amino acid insertion in a flexible loop (loop 1) that closes the active site after substrate binding (Söderholm et al. 2015). The three extra amino acids make the loop more flexible, allowing the positively charged guanidino group of R18 to stabilize binding of the negatively charged TrpF substrate (PRA) but simultaneously displaces a tryptophan (W145) that is necessary for binding the native substrate (Newton et al. 2017). Similarly, the L169R mutation placed an arginyl residue appropriately to fulfil the analogous role from a position below the active site, abolishing the binding of the native substrate (Newton et al. 2017). In the current experiments, all evolutionary paths involving HisA mutations started with a single amino acid substitution, Q18R, placing an arginine in the same position as the dup13-15 mutation but without loop expansion. The Q18R mutation, partially characterized in previous experiments, displays less rescue of Δ*trpF* growth than dup13-15, although it retains near wild-type HisA function (Lundin et al. 2020). This suggests the guanidino group of the arginine at position 18 in Q18R can reach a similar position as in the dup13-15 mutation, but the absence of extra amino acids in the loop may prevent the arginine from reaching its optimal position. Interestingly, three additional HisA mutations that improved the Δ*trpF* growth rescue of the Q18R mutant were found in the vicinity of position 18: H17R, A22T, and R23L. Each of these mutations improved overall growth in the evolution experiments but negatively impacted HisA function (Figure 3), suggesting the mutations enhanced the ability to substitute for TrpF *in vivo* at the expense of HisA function. If the normal-sized loop of the Q18R mutant is not flexible enough to allow the arginine to reach deeply into the active site for efficient PRA binding or catalysis, these other mutations may make the loop more flexible, allowing the arginine to reach its optimal position, potentially improving the ability to substitute for TrpF *in vivo*. Added flexibility due to the adjacent mutations could act analogously to the loop extension in the dup13-15 mutant, compromising the loop structure and negatively impacting native catalysis. Other mutations in HisA are more difficult to rationalize; V168 and F199 pack against each other in the hydrophobic core, and the mutations V168A and F199L likely destabilize the native structure, at least locally. Another mutation (A127G) is close to V168 and F199 but points into the barrel, forming part of the bottom of the active site cavity. These mutations may destabilize the structure, negatively impacting enzyme activity or steady-state levels, but making the active site more flexible in accommodating the alternative substrate. Other mutations in HisA affect surface-exposed charged or polar residues (T52, K107, R131), changing them to other charged or polar amino acids (T52S, K107E, R131H). Two of these (T52S, R131H) had modest or insignificant impacts on HisA function (Figure 3) while improving growth after tryptophan depletion, suggesting a different mode of action than the other mutations.

The evolution of bifunctional HisA is an example of substrate ambiguity, where an enzyme evolves to accommodate two different substrates while continuing to catalyze its original reaction. The main challenge is enabling the binding of the new substrate without losing the ability to bind the original one. For TrpA to retain its native function while evolving to substitute for TrpF, a case of catalytic promiscuity, seems less likely at first glance. Not only must it bind a new substrate and catalyze an entirely different reaction, but it must also do so without losing its original function, which appears to be a very difficult problem.

However, arguments can be made that the mutant TrpA variants characterized by Evran *et al*. (Evran et al. 2012) were already bifunctional. *In vivo*, TrpA depends on TrpB for activity (Pittard and Yang 2008), and as observed by us, this also applies to the detectable rescue of a *trpF* mutant. In their selections and growth rate measurements Evran *et al*. expressed the mutant *trpA* alleles from multicopy plasmids, leading to competition for TrpB by the two TrpA variants. If the expression of plasmid-borne TrpA was sufficiently high, the mutant TrpA would outcompete the wild-type TrpA. If the mutant TrpA variants lacked TrpA activity, rescuing a *trpF* mutant would not be possible. Alternatively, though perhaps less likely, to gain TrpF activity in their selection, TrpA may have first acquired mutations that prevented association with TrpB and allowed it to function efficiently without its partner.

#### Additional Mutations in Evolved Populations Potentially Affecting Fitness in the evolution experiments

Beyond the likely frameshift suppressor mutations identified in populations with the *trpA*(P62fs) mutation, many evolved lineages harboured additional mutations. These mutations could potentially influence growth in the conditions of the evolution experiment in several different ways.

Several populations contained mutations known to provide selective advantages in minimal glucose environments (e.g., *pykF*, *rpoS*, *spoT*, (King et al. 2004; Phillips et al. 2016; Knöppel et al. 2018). This suggests the evolved populations adapted not only to overcome the tryptophan deficiency but also for efficient growth or survival under the specific experimental conditions.

**Metabolic adjustments affecting the tryptophan or histidine pathways:** Some mutations might directly modify the impact of the causative mutations. These could allow for more growth without restoring tryptophan synthesis. Potential benefits include reduced wasteful consumption of precursors like PRPP and chorismate by the non-functional tryptophan pathway in a Δ*trpF* mutant, or potentially mitigating a potential toxic effect by the accumulation of the TrpF substrate. However, none of these potential benefits were experimentally verified through reconstruction and testing of the specific mutations.

Most populations displayed mutations affecting other genes within the *his* and *trp* operons. For instance, *hisG* mutations, potentially reducing the flow of PRPP into the histidine pathway, were identified in six out of ten sequenced non-mutators and four out of eight sequenced mutators from evolution experiment 2. Notably, these *hisG* mutations always co-occurred with *trpA* mutations (Table S1).

More than half of the whole-genome sequenced populations harboured mutations affecting TrpD and/or TrpE, seemingly independent of the presence or absence of TrpF-activating mutations. The nature of some of these mutations suggests negative impacts on protein function (e.g., frameshifts, a 98 bp deletion, premature stop codons). The 98 bp deletion removes the last 32 amino acids from TrpE and fuses it translationally to TrpD, most likely leading to a non-functional fusion protein. This mutation was found in a population that did not evolve to grow after tryptophan depletion and, if it results in the loss of functional TrpE and/or TrpD it may represent an evolutionary dead-end that has higher fitness than the ancestor but is unable to evolve further to allow growth after tryptophan depletion.

While other TrpD and TrpE mutations were also observed in populations that did not grow after tryptophan depletion, their impact on protein function is less clear. However, several of these mutations were present in populations with *hisA* and *trpA* mutations that allowed post-depletion growth, suggesting that they still allow for some level of TrpD and TrpE protein production. Among these, frameshifts and nonsense mutations were predominant among the non-mutators while the mutator populations were dominated by single amino acid substitutions. This pattern mirrors the differences in mutation spectra observed in *hisA* and *trpA* between mutator and non-mutator populations, and may have the same explanation: in the non-mutator, the rate of amino acid substitutions is orders of magnitude lower than the rate of frameshift mutations, leading to an apparent bias for frameshift mutations among the selected mutations.

Given the requirement for functional TrpD and TrpE for growth after tryptophan depletion, the frameshifts and nonsense mutations are likely to be inherently leaky through translational frameshifting and stop codon readthrough. Loss or reduced expression or activity of these enzymes could potentially limit the flow of precursors into the tryptophan pathway, impacting growth or viability of the Δ*trpF* mutant under tryptophan-limiting conditions.

Two populations with *hisA* mutations displayed mutations in *hisH*, the preceding gene in the *his* operon. As the specific mutations are amino acid substitutions located upstream of the *hisA* start codon, they could potentially affect HisH function or even influence *hisA* translation initiation, potentially increasing HisA expression (leading to improved growth rescue).

The evolution experiments were conducted in a medium lacking guanosine. Since our later findings revealed that *hisA* mutations could be rescued by guanosine supplementation, it's possible that any mutations affecting flux through the histidine or tryptophan pathways (*hisG*, *trpD*, *trpE*, and possibly *hisH*) might act through effects on the PRPP pool, and that some of these would not have been seen if the evolution experiments were done in medium with added guanosine.

**Regulation of Histidine Biosynthesis:** One population (DA65453) contained a mutation affecting $tRNA^{His}$ (the *hisR* gene), and two additional populations (DA59062, DA59064) had different mutations in the promoter of the *argX*-*hisR*-*leuT*-*proM* tRNA operon that likely reduce the expression of the tRNAs. Reduced expression or function of $tRNA^{His}$ is expected to lead to over-expression of the *his*-operon by affecting the regulation of transcription attenuation in the *his*-operon leader region, mimicking histidine starvation (Winkler and Ramos-Montañez 2009). The mutations affecting *hisR* are thus likely to cause increased expression of HisA (and thus increase expression of the evolving growth rescue function).

### Supplementary Figures

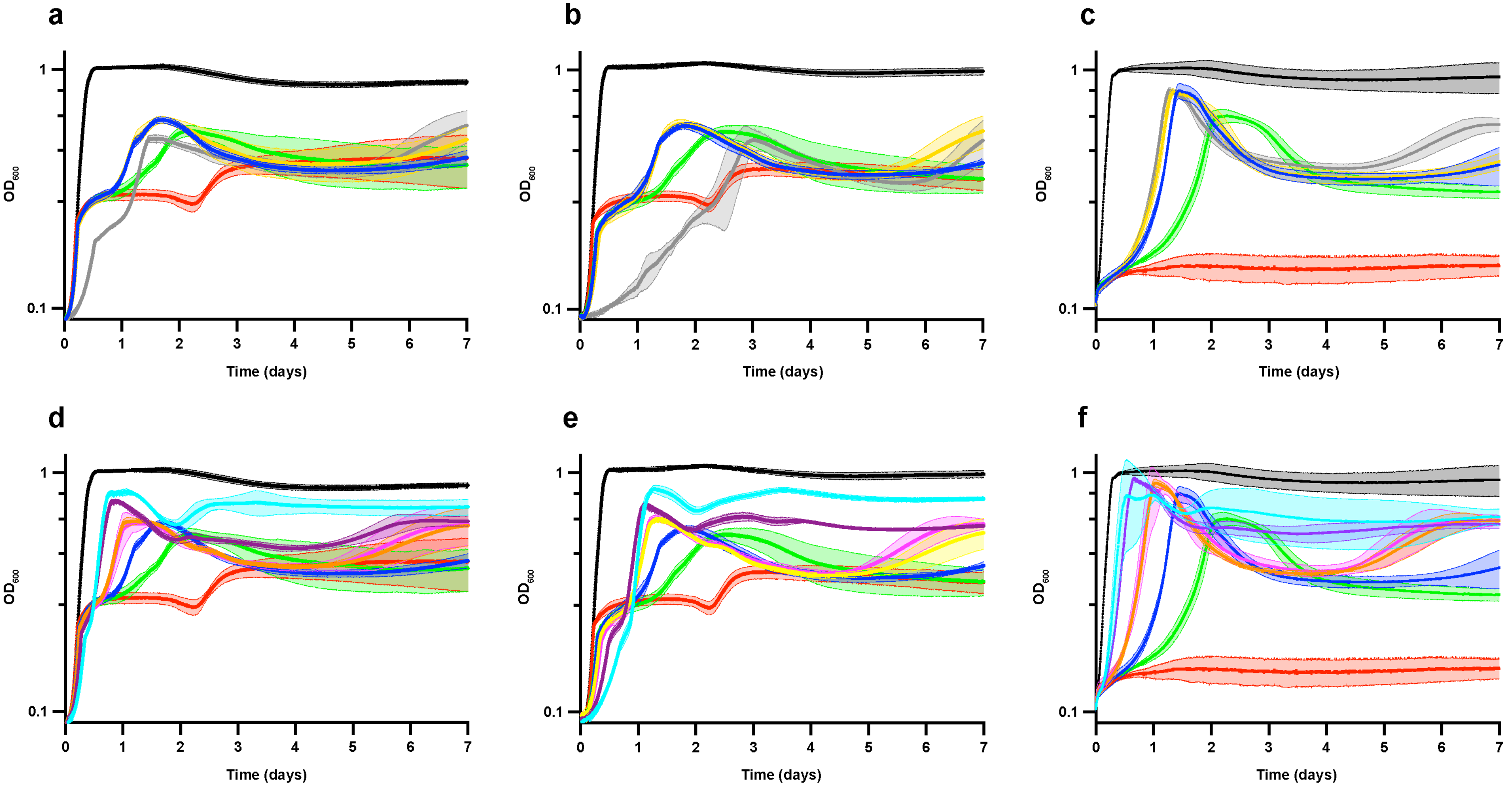


**Figure S1.** Growth of re-constructed *hisA* mutants. In all graphs the ancestral (*ΔtrpF*; red) and a wild-type (*trpF*+; black) strain is included for comparison. The curves show the average $OD_{600}$ with standard deviation of at least four biological replicates. **(a, d)** Cultures grown after 1000x dilution in 2x M9 + 0.4 % glucose supplemented with 5 µM tryptophan + 3 mM guanosine. **(b, e)** Cultures grown after 1000x dilution in 2x M9 + 0.4 % glucose supplemented with 5 µM tryptophan (no guanosine). **(c, f)** Cultures grown after 100x dilution in 2x M9 + 0.4 % glucose without supplementation. **(a - c)** *hisA* mutations found at 120 and 170 generations in population 1-1. Green; *hisA*(Q18R), grey; *hisA*(H17R Q18R), yellow; *hisA*(Q18R A22T), blue; *hisA*(Q18R K107E). **(d - f)** *hisA* mutations found at 320 and 420 generations in population 1-1. Green; *hisA*(Q18R; included for comparison), blue; *hisA*(Q18R K107E), pink; *hisA*(Q18R I62M K107E); orange; *hisA*(Q18R K107E V168A), purple; *hisA*(Q18R R23L K107E V168A), cyan; *hisA*(Q18R A22T R23L K107E V168A).

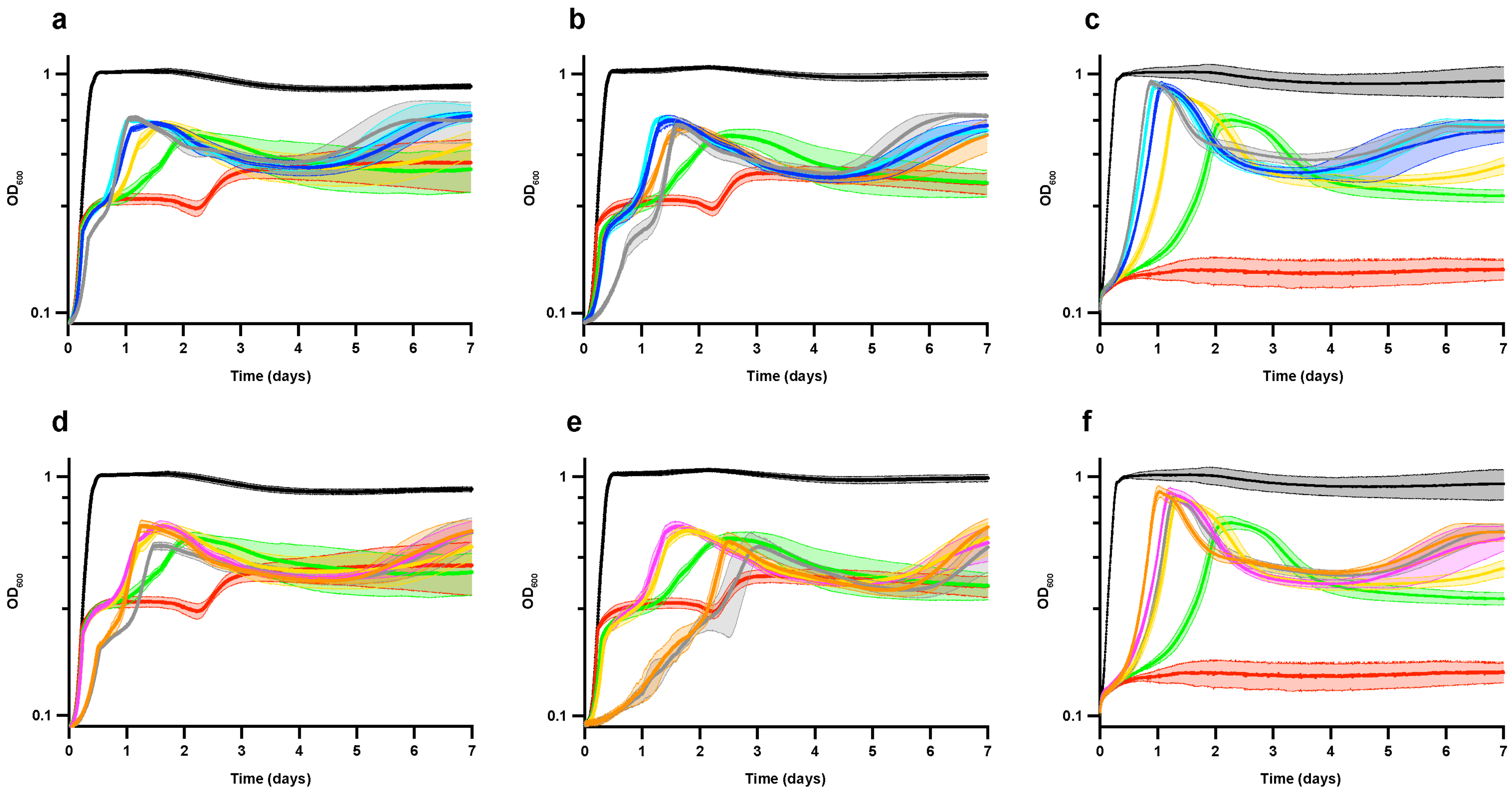


**Figure S2.** Growth of re-constructed *hisA* mutants, continued. In all graphs the ancestral (*ΔtrpF*; red) and a wild-type (*trpF*+; black) strain is included for comparison. The curves show the average $OD_{600}$ with standard deviation of at least four biological replicates **(a, d)** Cultures grown after 1000x dilution in 2x M9 + 0.4 % glucose supplemented with 5 µM tryptophan + 3 mM guanosine. **(b, e)** Cultures grown after 1000x dilution in 2x M9 + 0.4 % glucose supplemented with 5 µM tryptophan (no guanosine). **(c, f)** Cultures grown after 1000x dilution in 2x M9 + 0.4 % glucose without supplementation. **(a - c)** *hisA* mutations found in population 1-3. Green; *hisA*(Q18R), yellow; *hisA*(Q18R A22T), cyan; *hisA*(Q18R A22T V168A), grey; *hisA*(Q18R A22T I62M R131H), blue; *hisA*(Q18R A22T V168A F199L). **(d - f)** *hisA* mutations found in the *mutS* populations. Green; *hisA*(Q18R), yellow; *hisA*(Q18R A22T), orange; *hisA*(Q18R A127G), grey; *hisA*(H17R Q18R), pink; *hisA*(Q18R T52S).

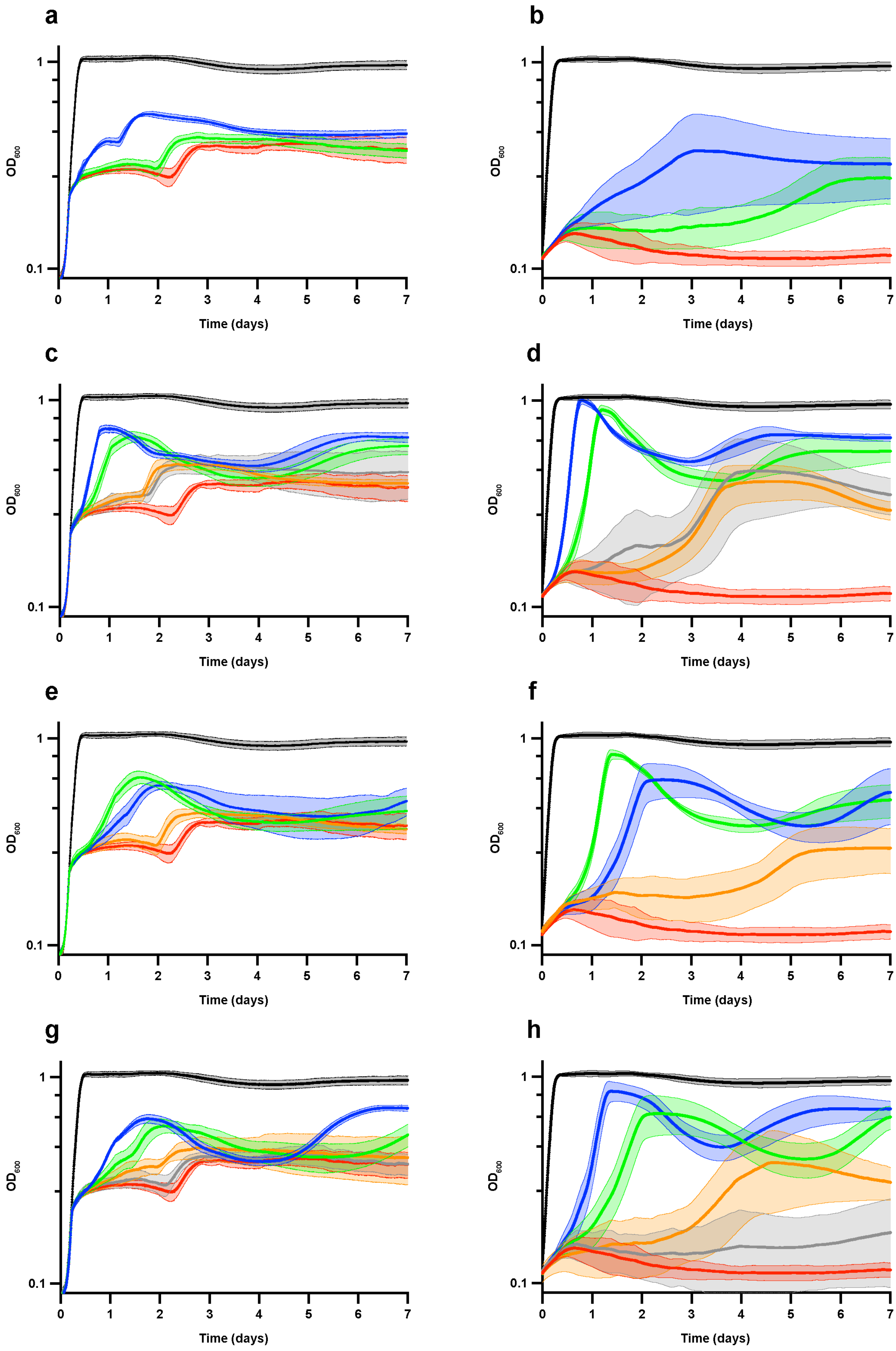

a
b
c
d
e
f
g
h
OD600
Time (days)
1
0.1
0
1
2
3
4
5
6
7

**Figure S3.** Growth of re-constructed *trpA* mutants. In all graphs the ancestral (*ΔtrpF*; red) and a wild-type (*trpF*+; black) strain is included for comparison. The curves show the average $OD_{600}$ with standard deviation of five biological replicates. **(a, c, e, g)** Cultures grown after a 1000x dilution in 2x M9 + 0.4% glucose supplemented with 5 µM tryptophan + 3 mM guanosine. **(b, d, f, h)** Cultures grown after a 100x dilution in 2x M9 + 0.4% glucose without supplementation. **(a, b)** *trpA* mutations found in the non-mutator populations. Green; *trpA*(A67T), blue; *trpA*(P62fs). **(c, d)** *trpA* mutations found in mutator populations 2*mutS*4, 2*mutS*11, and 2*mutS*23. Grey; *trpA*(D27Y), orange; *trpA*(G98C), green; *trpA*(D27Y G98S), blue; *trpA*(D27Y G98C). (e, f) *trpA* mutations found in mutator population 2*mutS*14. Orange; *trpA*(Y102H), blue; *trpA*(G98S Y102H), green; *trpA*(G98S Y102H Q250R). (g, h) *trpA* mutations found in mutator population 2*mutS*16. Grey; *trpA*(T24S), orange; *trpA*(G61S), green; *trpA*(T24S G61S), blue; *trpA*(T24S G61S P62S).

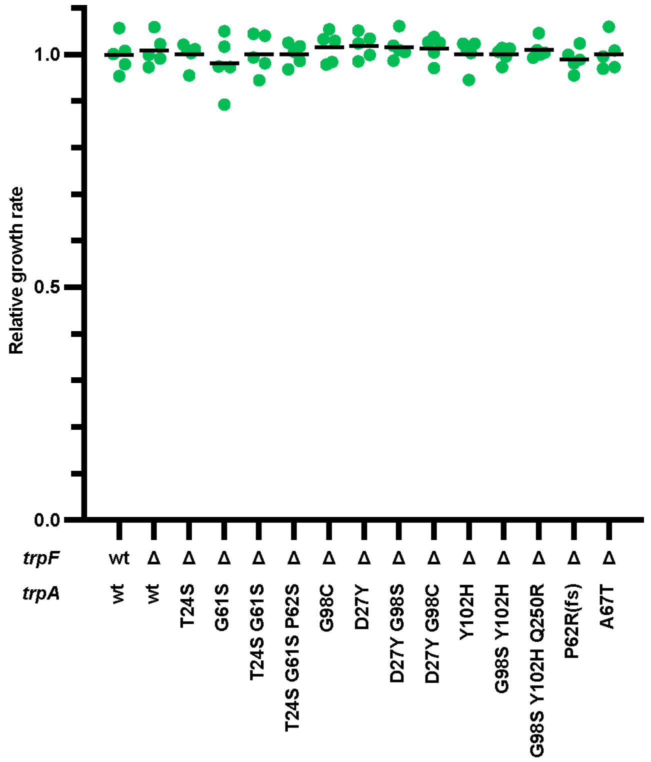


**Figure S4.** Growth rates of *trpA* mutants prior to tryptophan depletion. Exponential growth rates were determined in the early rapid growth phase (0.095 > $OD_{600}$ < 0.184) in M9 glucose medium supplemented with tryptophan (5 µM) and guanosine (3 mM), and set relative to a *trpF*(wt) *trpA*(wt) strain grown in the same experiment. All strains were grown as five biological replicates, and none of the strains grew differently from the *trpF*(wt) *trpA*(wt) strain according to a two-tailed Student's t-test with equal variance.

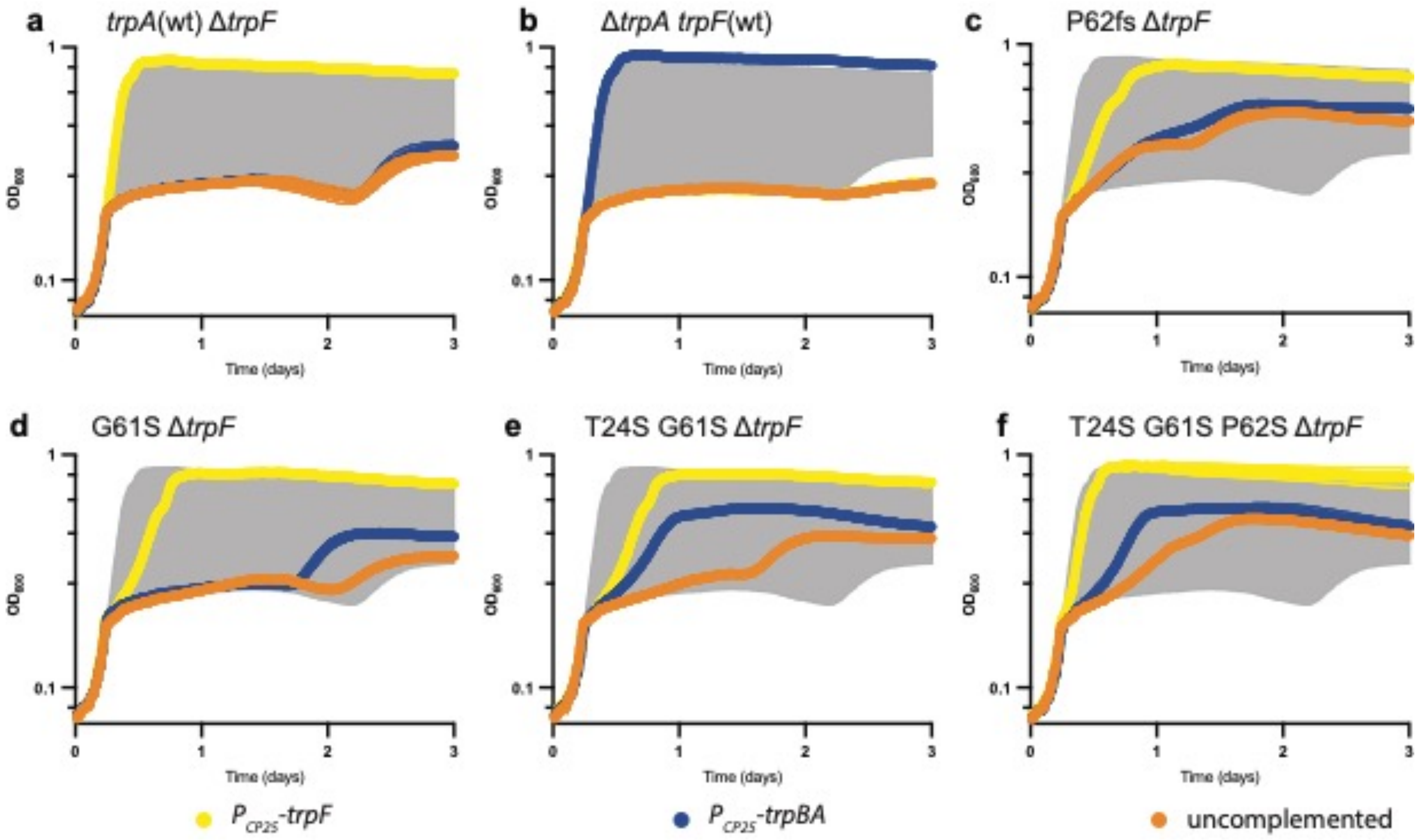


**Figure S5.** Growth of complemented *trpA* mutants. The curves show the average $OD_{600}$ with standard deviation of five biological replicates (most of the error ranges are too small to be seen). To reduce noise from a faulty bioscreen machine, the data was smoothed by taking the median in an eleven data point sliding window. All cultures were grown after a 1000x dilution in 2x M9 + 0.4% glucose supplemented with 5 µM tryptophan. **(a)** *trpA*+ Δ*trpF* (ancestor for the evolution experiments) **(b)** Δ*trpA trpF+* **(c)** *trpA*[P62fs] Δ*trpF* **(d)** *trpA*[G61S] Δ*trpF* **(e)** *trpA*[T24S G61S] Δ*trpF* **(f)** *trpA*[T24S G61S P62S] Δ*trpF*. Orange; uncomplemented strains, blue; strains complemented with *trpBA*; yellow; strains complemented with *trpF.* The grey shaded area shows the surface between the curves of the *trpA*(wt) Δ*trpF* strain without complementation and complemented with *trpF* (orange and yellow curves in (a)).

**a**

*trpA*(wt)

```
              60 61 62
UCCGAUCCGCUGGCCGAUGGCCCUACCAUCCAGAAUGCGAACUUACGCGCCUUCGCCGCUGGCGUCACGCCGGCUCAGUGUUUUGAAAUG
 S  D  P  L  A  D  G  P  T  I  Q  N  A  N  L  R  A  F  A  A  G  V  T  P  A  Q  C  F  E  M
```

**b**

*trpA*(P62fs)

```
              60 61 62
UCCGAUCCGCUGGCCGAUGGCCGAUGGCCCUACCAUCCAGAAUGCGAACUUACGCGCCUUCGCCGCUGGCGUCACGCCGGCUCAGUGUUUUGAAAUG
 S  D  P  L  A  D  G  R  W  P  Y  H  P  E  C  E  L  T  R  L  R  R  W  R  H  A  G  S  V  F  *
  P  I  R  W  P  M  A  D  G  P  T  I  Q  N  A  N  L  R  A  F  A  A  G  V  T  P  A  Q  C  F  E  M
```

**Figure S6.** A frameshift mutation in *trpA* generates a bifunctional gene. **(a)** Codons 55 – 84 in wild-type *trpA*. Top, mRNA sequence; bottom, amino acid sequence. Bold text: five nucleotide repeats (UGGCC) involved in generation of the P62fs mutant. Blue box: The 7 bp sequence (UGGCCGA) that is duplicated in the P62fs mutant. **(b)** The corresponding sequence in the *trpA*(P62fs) mutant. Red text in amino acid sequence: mis-incorporated amino acids if translation continues in the original (zero) reading frame. Red text in mRNA: A UGA stop codon encountered in the zero frame in the Pro62fs mutant. Blue text in amino acid sequence: mis-incorporated amino acids if translation shifts into the +1 frame. The amino acid at position 60 (D60) is one of the catalytic residues for the native reaction, probably limiting the window of functional suppression to somewhere after D60 and before the zero frame stop codon.

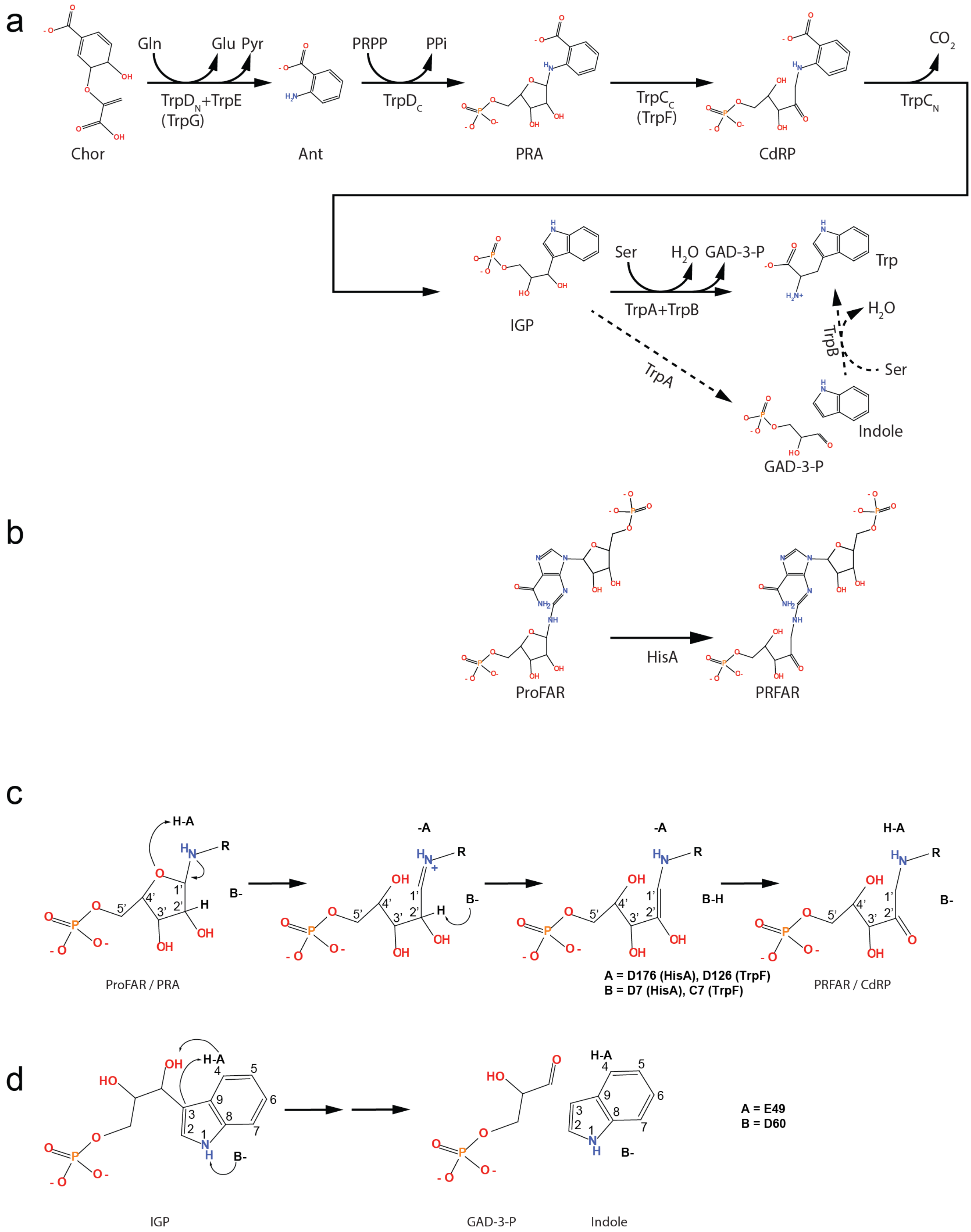


**Figure S7. (a)** Biosynthesis of tryptophan is catalyzed by the enzymes indicated below the reaction arrows. Subscript letters (N or C) refer to which domain of a fused protein (in *Salmonella* and other members of *Enterobacteriaceae*) contain the indicated activity, and protein names in parenthesis are the names of the corresponding proteins in organisms where the enzymes are not fused. For simplicity, we refer to the enzyme PRA isomerase contained within the C-terminal domain of TrpC as TrpF, despite it not being a separate protein in *Salmonella*. Gln, L-glutamine; Glu, L-glutamate; PRPP, phosphoribosyl-pyrophosphate; PPi, inorganic phosphate; $CO_2$, carbon dioxide; Ser, L-serine; GAD-3-P, glyceraldehyde-3-phosphate; Chor, chorismate; Ant, anthranilate; PRA, phosphoribosyl anthranilate; CdRP, 1-(2-carboxyphenylamino)-1’-deoxyribulose-5’-

phosphate; IGP, indole-3-glycerol phosphate; Trp, L-tryptophan. Dashed reaction arrows: while the physiological activity of tryptophan synthase (TrpA+TrpB) is the direct conversion of IGP to Trp (without the release of Indole as an intermediate), *in vitro* the reaction can be broken up into two separate reactions. **(b)** HisA (ProFAR isomerase) catalyzes the isomerization of N'-[(5'-phosphoribosyl)-formimino]-5-aminoimidazole-4-carboxamide ribonucleotide (ProFAR) to N'-[(5'-phosphoribulosyl) formimino]-5-aminoimidazole-4-carboxamide-ribonucleotide (PRFAR), the fourth step in histidine biosynthesis. **(c – d)** Part of the catalytic mechanisms for HisA, TrpF and TrpA. **(c)** The reactions catalyzed by HisA and TrpF. The general acid (D176 in HisA, D126 in TrpF) protonates the ribose ring oxygen, and the general base (D7 in HisA, C7 in TrpF) abstracts a proton from the C2' of the ribose. **(d)** The reaction catalyzed by TrpA. The general acid (D60) protonates the C3 carbon, and the general base (E49) abstracts a proton from the N1 nitrogen of the indole.

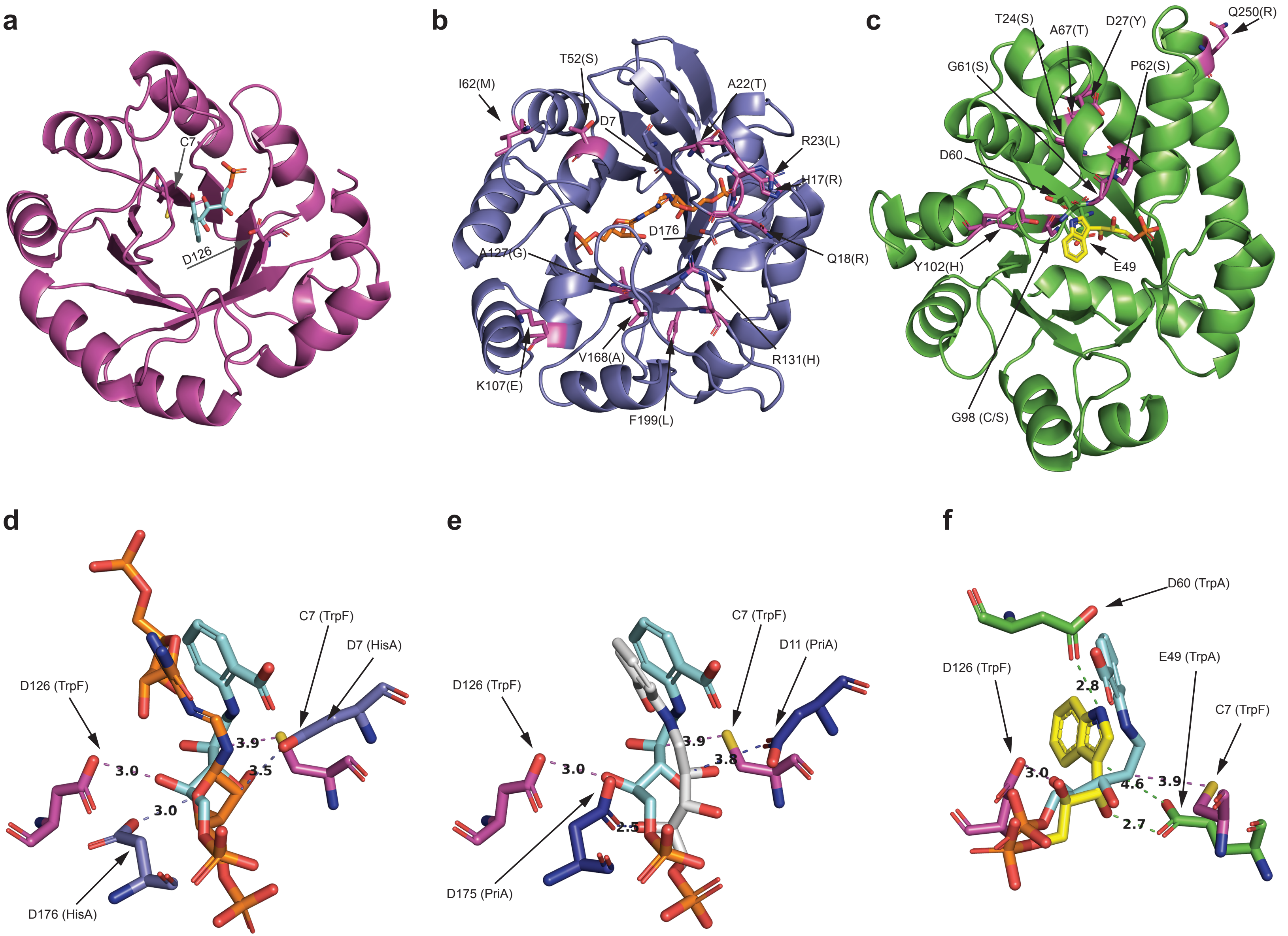

a
C7
D126
b
I62(M)
T52(S)
D7
A22(T)
R23(L)
H17(R)
D176
Q18(R)
A127(G)
V168(A)
R131(H)
K107(E)
F199(L)
c
T24(S)
A67(T)
D27(Y)
Q250(R)
G61(S)
P62(S)
D60
Y102(H)
E49
G98 (C/S)
d
C7 (TrpF)
D7 (HisA)
D126 (TrpF)
3.9
3.0
3.5
3.0
D176 (HisA)
e
C7 (TrpF)
D11 (PriA)
D126 (TrpF)
3.9
3.0
3.8
2.5
D175 (PriA)
f
D60 (TrpA)
D126 (TrpF)
E49 (TrpA)
C7 (TrpF)
2.8
3.0
4.6
3.9
2.7

**Figure S8.** Crystal structures of **(a)** *Thermotoga maritima* TrpF (pdb code: 1lbm) with bound product analogue rCdRP (cyan C-atoms), **(b)** *S. enterica* HisA (D7N D176A; pdb code: 5a5w) with bound substrate ProFAR (orange C-atoms), *in silico*-reverted to wild-type sequence to show the likely positions of the general acid and base, and **(c)** *S. enterica* TrpA (D60N; pdb code: 1a5b) with bound substrate IGP (yellow C-atoms), *in silico*-reverted to wild-type sequence. **(d)** Superposed HisA and TrpF, showing only the locations of the ligands and the general acids and bases, and the distances between the catalytic residues and reacting atoms. **(e)** Same as in (d) but TrpF and *Mycobacterium tuberculosis* PriA (pdb code: 2y85) with bound substrate analogue rCdRP (residues from PriA with dark blue C-atoms, and rCdRP with light grey C-atoms). **(f)** Same as (d) but TrpA and TrpF. The locations of mutations in HisA (b) and TrpA (c) are shown with magenta C-atoms and magenta cartoon. Arrows point at catalytic residues and mutant positions, with wild-type residues before parentheses and mutant residues within parentheses. The images and *in silico* mutagenesis were made with PyMOL (Schrödinger, LLC 2021). PriA and HisA were superposed in PyMOL using ‘align’ (RMSD = 1.335 over 1003 atoms), TrpA and TrpF were superposed on HisA using ‘cealign’ (RMSD = 4.888 and 4.593 over 176 residues for TrpF:HisA and TrpA:HisA, respectively).
